# Evolution of the ribosomal exit tunnel through the eyes of the nascent chain

**DOI:** 10.64898/2025.12.25.696519

**Authors:** Tomasz Włodarski

**Author notes:** Correspondence: TW.

## Abstract

The exit tunnel is a universally conserved feature of the ribosomal large subunit that directs the nascent polypeptide chain into the cellular environment and is involved in co-translational folding, stalling, and antibiotic binding. While recent advances in cryo-EM have revealed organism-specific variations in ribosome structure, tunnel definition and comparative analyses have largely relied on pure geometric algorithms that often approximate the tunnel as a static, single tube with varying radius. Here, we present a functional, nascent-chain-centric characterisation of the exit tunnel across the tree of life, derived from all-atom molecular dynamics simulations using the most complete set of 55 distinct cytoplasmic ribosome structures. By mapping the steric accessibility through the “eyes” of the nascent chain (NC) at five different stages of translation, we reveal a topological and stage-dependent complexity invisible to static geometric approaches, demonstrating how tunnel accessibility dynamically changes during biosynthesis. We identify transiently accessible, bacteria-specific lateral branches (“side tunnels”) that effectively expand the functional volume of the tunnel but may also serve as kinetic traps for the nascent chain. In contrast, we show that archaeal and eukaryotic ribosomes have structurally occluded these cavities by incorporating the eL39 protein. These evolutionary “plugs” seal the tunnel wall and decrease its functional width. Collectively, our results demonstrate that the ribosome exit tunnel is not a simple linear tube, but a branched, lineage-specific landscape where accessibility is temporally gated by nascent chain length. This functional definition provides a new framework for understanding how ribosomal architecture modulates the early stages of protein biogenesis.

## Introduction

The ribosome is an ancient biomolecular machine responsible for translating genetic information into proteins in all living systems (1). After peptide bond formation in the peptidyl transfer centre (PTC) (2), the nascent polypeptide exits the ribosome through a narrow channel - the ribosomal exit tunnel. This structural feature is conserved across all domains of life (1); remarkably, every protein ever produced by life on Earth has passed through it.

The tunnel, including its vestibule, is universally constructed from ribosomal RNA and conserved ribosomal proteins, including uL4, uL22, uL23, uL24, and uL29. Eukaryotes and archaea also incorporate the specific protein eL39, which replaces the uL23 exit tunnel loop. This tunnel provides the first opportunity for the nascent chain (NC) to fold; however, due to its narrow diameter, only α-helices and small β-hairpins can form within it (3–9). More complex structures fold in or beyond the vestibule as part of the co-translational folding process (7, 10–18). Modifications to the exit tunnel can directly influence the co-translational folding, either through rational engineering of tunnel loops (19, 20) or by using ribosomal protein paralogs, such as RPL39L (21). Additionally, the tunnel is a regulatory hotspot: it interacts with specific nascent chains to modulate elongation and induce translational stalling (22–24), and it is a well-established target for ribosome-binding antibiotics (25, 26).

Historically, the existence of the tunnel was inferred from proteolytic experiments (27), confirmed with antibody binding (28), and finally visualised by electron microscopy (29). Earlier low-resolution cryo-EM studies suggested the presence of possible side branches and alternative exits (30). However, the first in-depth analysis of the ribosome tunnel, conducted using the *H. marismortui* ribosome with a rolling ball algorithm (31), found no tunnel branching and confirmed a narrow geometry that precludes substantial folding.

Subsequent molecular dynamics (MD) simulations further characterised the physicochemical complexity of the tunnel using the *H. marismortui* ribosome. The 3D potentials of mean force (PMFs) calculated for five representative amino acids showed that distinct residues experience different energetic landscapes throughout the ribosomal exit tunnel (32). As each 3D PMF differed significantly, this suggests the tunnel is sensitive to the chemical nature of the nascent chain. Mapping the electrostatic potential has also shown that the tunnel is highly negatively charged and heterogeneous (33), which can modulate elongation speed (34, 35). Furthermore, water dynamics within the tunnel differ significantly from bulk water (36), potentially weakening the hydrophobic effect in the vestibule region (37), highlighting the complexity of this confined environment.

A comparative study of 20 distinct ribosome structures across all domains of life recently revealed that tunnel geometry broadly mirrors phylogenetic relationships and identified a previously unrecognised “second constriction” site in eukaryotes (38). It also found that the upper part of the tunnel is generally more structurally conserved than the lower part. Subsequently, it has been proposed that the resulting narrower eukaryotic tunnel could act as a kinetic trap for small proteins (39). Most recently, this geometric classification has been extended and refined to show that certain eukaryotic lineages, such as parasitic protists, can revert to wider, bacteria-like tunnel architectures (40). Additionally, we recently demonstrated that the tunnel vestibule may support at least two main NC pathways, with a preference that depends on folding status and translation stage (41).

Despite these significant insights, most previous studies have focused on overall geometry (radius and volume) rather than on how the nascent chain experiences the tunnel, especially across different stages of its biosynthesis. Here, we adopt a functional approach to explore the structural heterogeneity of ribosome tunnels across various species by analysing it through the “eyes” of the nascent chain. To map the shape of the tunnel, we followed all-atom MD trajectories of the NC and converted them into 3D density maps that reflect the spatial occupancy of the nascent chain within the tunnel. Moreover, we generated these maps for different stages of translation, based on the hypothesis that the local accessibility of the tunnel changes during biosynthesis.

We leverage the recent increase in cryo-EM ribosome structures, which has almost tripled the number of unique high-resolution cytoplasmic ribosomes from 18 to 55 (38). Our structural analysis confirms the previously identified geometric constriction sites but reveals a far more complex topology defined by transient accessibility. We characterised a third constriction site defined by eukaryotic/archeal-specific eL39 protein and identified additional bifurcation tunnels (“side tunnels”) observed exclusively in bacteria, which are gated by nascent chain length. Interestingly, in eukaryotes and archaea, these branches are closed by alternative conformations of conserved proteins or rRNA, specifically through the incorporation of the eL39. These additional tunnel branches are only accessible to the NC of certain lengths and provide a compelling representation of how the tunnel architecture is experienced differently by the NC at early stages of translation. Using this functional approach, we demonstrate that the tunnel cannot be accurately described as a simple linear cylinder with a varying radius, as its topology is far more complex. Finally, we mapped the changes in the physicochemical environment along the tunnel pathway, providing a rich resource for future studies of NC-tunnel interactions and the discovery of potential drug-binding sites.

## Results

### Comprehensive identification of ribosome structures across the tree of life

Recent advances in cryogenic electron microscopy (cryo-EM) (42, 43) have led to a significant increase in high-resolution ribosome structures, enabling detailed comparative analysis. To systematically explore the structural diversity of ribosomal exit tunnels across the tree of life, we conducted a comprehensive search of the Protein Data Bank (PDB) (44) (see Methods). This search identified 68 unique high-resolution structures: 18 bacterial, 45 eukaryotic (including 11 mitochondrial and one chloroplast ribosome), and five archaeal (Table S1-S4). For the present study, we focused on the 56 cytoplasmic ribosomes, as organellar ribosomes, particularly mitochondrial ones, exhibit highly divergent tunnel architectures and were therefore excluded from the analysis. The ribosome from *Cutibacterium acnes* was also omitted due to the presence of a bound antibiotic in the exit tunnel, which would interfere with structural analysis, leaving us with a final set of 55 ribosomes.

### Tunnel alignment and PTC structural conservation across species

To enable direct structural comparisons of ribosomal exit tunnels, we aligned all selected ribosome structures to the *E. coli* reference tunnel (PDB ID: 7zp8 (19)) and extracted the corresponding tunnel regions (Fig. 1a; see Methods). Analysis of the PTC, which is the most conserved region of the tunnel, confirmed the high quality of this structural alignment, with average backbone root-mean-square deviations (RMSDs) of < 2 Å relative to *E. coli* for nearly all structures (Fig. S1a, c). Despite this high structural similarity, which lies well within the resolution limits of the cryo-EM maps, clustering analysis broadly recapitulates established phylogenetic relationships (Fig. S1b). A notable exception was the archaeal ribosome from *Thermococcus kodakarensis*, which displayed a higher RMSD of 3.24 Å (Fig. S1c), primarily due to an unusual 23S rRNA conformation (nucleotides 2170-2173; see below).

**Figure 1.**
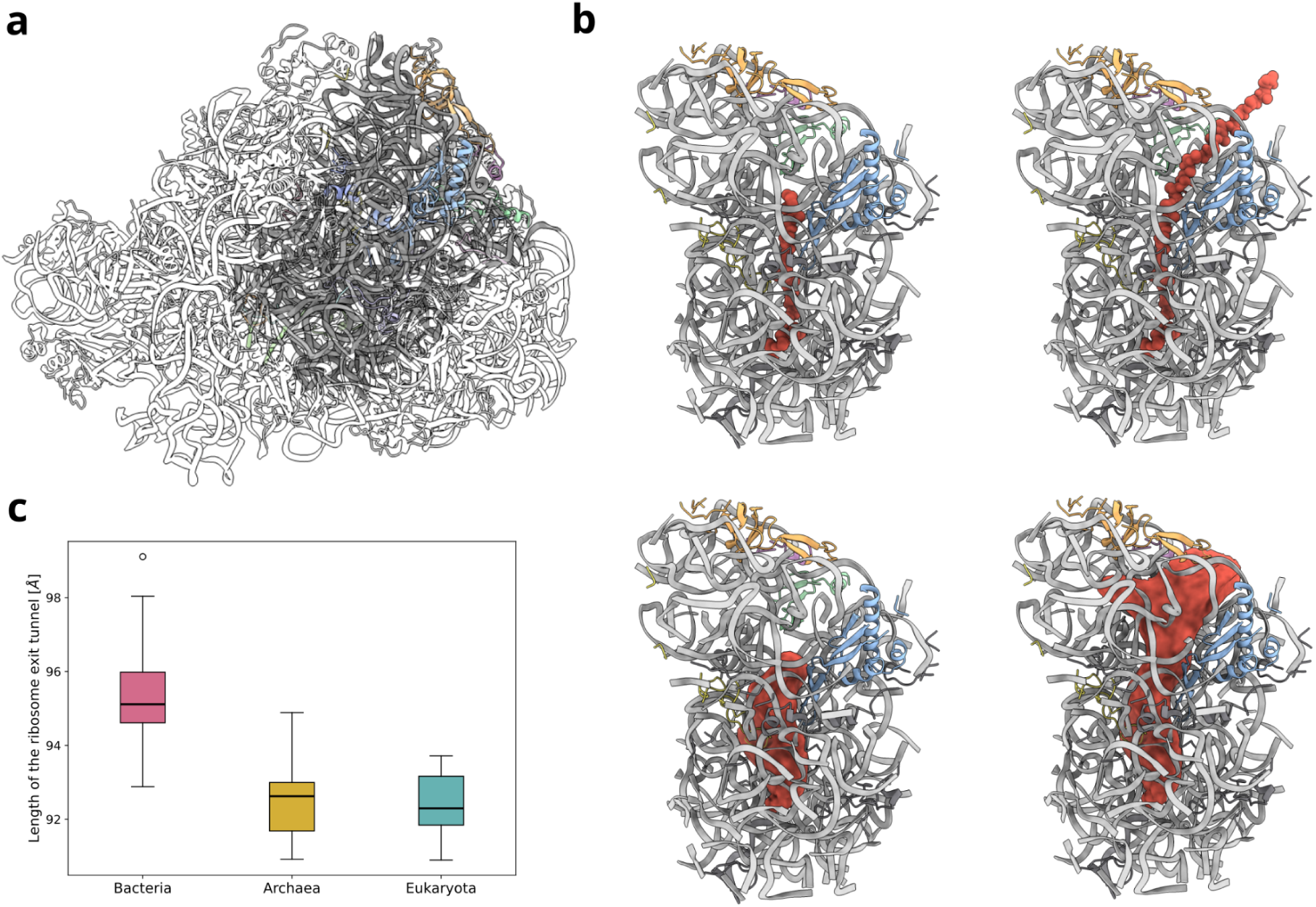
Structural definition of the ribosomal exit tunnel. **a)** The *E. coli* 50S ribosome structure (PDB ID: 7zp8 (19)), highlighting the region selected as the reference exit tunnel. **b)** Visualisation of the functional tunnel definition. **Top:** The representative snapshots of a nascent chain (red atoms) within the exit tunnel at two different stages of biosynthesis. **Bottom:** The corresponding 3D occupancy maps (red volume) derived from MD trajectories, capturing the functional volume accessible to the nascent chain. Ribosomal proteins are colored: uL4 (yellow), uL22 (blue), uL23 (green), uL24 (orange), and uL29 (violet). **c)** Comparison of ribosomal exit tunnel lengths across the three main domains of life.

### A functional definition reveals conserved tunnel lengths across domains

To define the ribosomal exit tunnel in a functionally relevant manner, we adopted a nascent chain-centric approach that captures the accessible volume and boundaries of the tunnel based on NC dynamics, moving beyond the static geometric criteria used in previous studies (38, 40) (Fig. 1b, Fig. S2). We hypothesised that tunnel accessibility is dynamic: in the early stages of biosynthesis, distal regions of the tunnel are unreachable to the NC, whereas longer chains become entropically restricted from re-entering deeper segments. To capture this progressive evolution of accessible space, we generated five ribosome nascent chain complexes (RNCs) with chains of increasing lengths (10, 20, 30, 40 and 60 residues) (Fig. S2).

By subjecting these complexes to all-atom structure-based model (SBM) molecular dynamics simulations, we calculated average NC occupancy maps that define the functional shape of the tunnel - effectively providing a “plaster cast” of the NC-accessible space (Fig. 1b, Fig. S2). Based on these maps, we defined the main tunnel path and used it to quantify changes in tunnel dimensions (length, cross-sectional area, and asphericity) and physicochemical properties along the axis.

Establishing a consistent tunnel boundary across species presents a specific challenge: while the entrance is clearly marked by the 3’ end of the P-site tRNA, the exit opens gradually into the wider vestibule. We selected the plane of the uL24 loop, which prominently overhangs the vestibule in bacteria, as this universal marker (Fig. S3). Using this definition, we measured the length of each tunnel along its main path (Fig. 1c). In line with previous findings (38), bacterial exit tunnels are slightly longer (mean length: 95.5 ± 1.5 Å) than their eukaryotic (92.4 ± 0.7 Å) and archaeal (92.6 ± 1.3 Å) counterparts. This difference arises primarily from the extended uL24 loop in bacteria, which redirects the NC path and causes it to bend within the vestibule (Fig. S4a), whereas the core tunnel path remains highly conserved across all domains (Fig. S4b).

### Structural features of bacterial ribosome tunnels

We characterised the tunnel landscape for all 17 bacterial ribosome tunnels spanning seven phyla (Pseudomonadota, Spirochaetota, Bacillota, Actinomycetota, Deinococcota, Bacteroidota and Mycoplasmatota), providing broad phylogenetic coverage (Fig. S5). Despite this diversity, we observe that bacterial tunnels share a highly conserved core architecture with distinct, transiently accessible features (Fig. 2a, Fig. S6). From the perspective of the NC, the first major structural hurdle is a well-characterised constriction site located ∼30 Å from the PTC (Fig. 2a, d). Formed by the extended loops of uL4 and uL22, this narrowing is universal in bacteria, consistent with previous geometric analyses (38). However, we identified significant local divergence in this region: ribosomes from *T. thermophilus, M. smegmatis,* and *E. faecalis* display a markedly tighter constriction at ∼27 Å from the PTC (Fig. S5, S6a). This distinct narrowing arises from alternative conformations of residues corresponding to *E. coli* Arg61 (uL4) and U790 (23S rRNA), which sterically occlude this exit tunnel region (Fig. S7a). Whether this is a species-specific feature or reflects local conformational dynamics remains unknown, but it highlights how even subtle side-chain rearrangements can drastically modulate the effective geometry experienced by the nascent chain. A second constriction is located ∼45 Å from the PTC (Fig. 2a), defined by uL22 residues and surrounding rRNA (Fig. S7b). While this aligns positionally with the second constriction site observed in eukaryotes and archaea (∼50 Å from the PTC; see below), it remains significantly wider in bacteria.

**Figure 2.**
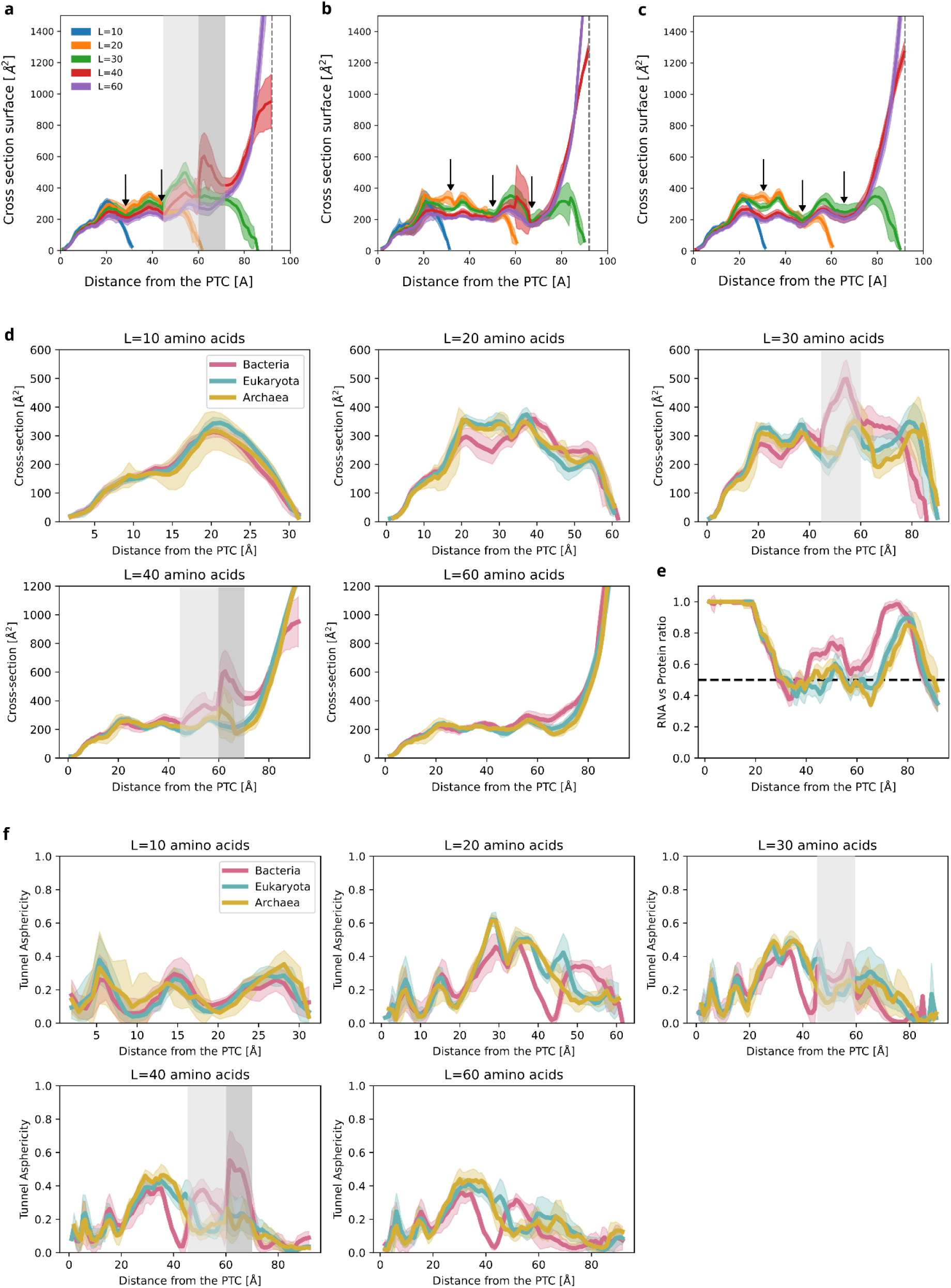
Evolutionary divergence of ribosomal exit tunnel geometry. **a-c)** Functional cross-sectional area profiles for **(a)** bacteria**, (b)** archaea and **(c)** eukaryotes, derived from MD simulations of nascent chains with increasing length (L=10 to 60 amino acids). Colour-shaded regions represent the spread across species, whereas the light and dark grey vertical regions represent the first- and second-side tunnels, respectively. **(d)** Cross-domain comparison of tunnel profiles for each NC length, highlighting conserved constriction sites and lineage-specific differences. **(e)** Tunnel wall composition mapped along the path, plotted as the ratio of rRNA to ribosomal protein. **(f)** Asphericity profiles quantifying the deviation from circular geometry (0 = circular; 1 = highly elongated), revealing domain-specific topological irregularities.

Beyond these conserved constrictions, our functional mapping revealed that tunnel accessibility is strictly modulated by nascent chain length, uncovering two previously underappreciated lateral branches or “side tunnels“ (Fig. 2a, d). First, we identified a proximal side tunnel 45 - 60 Å from the PTC, located beneath the extended loop of the uL23 protein (Fig. 2a and 3a). Although this branch is “closed” (it does not breach the ribosome surface), it acts as a significant volumetric expansion. Crucially, this region is particularly accessible to NCs of 30 residues but becomes less available as the chain lengthens, though it remains partially accessible at 40 residues in some species (Fig. S8). Access to this side tunnel appears to be gated by the side chains neighbouring rRNA and uL23 residues; for example, in *M. tuberculosis*, the protrusion of Arg66, Arg68, and Tyr73 restricts the NC entry (Fig. S8), suggesting a species-specific modulation of accessible space. Second, we observed a larger, distal side tunnel 60-70 Å from the PTC, opening between ribosomal helices H7 and H24 (Fig. 2a and 3b). This region becomes accessible at an NC length of 40 residues and is a conserved feature across all examined bacteria, with the notable exception of *L. monocytogenes*, where a tighter packing of helices H7 and H24 occludes the opening (Fig. S9).

**Figure 3.**
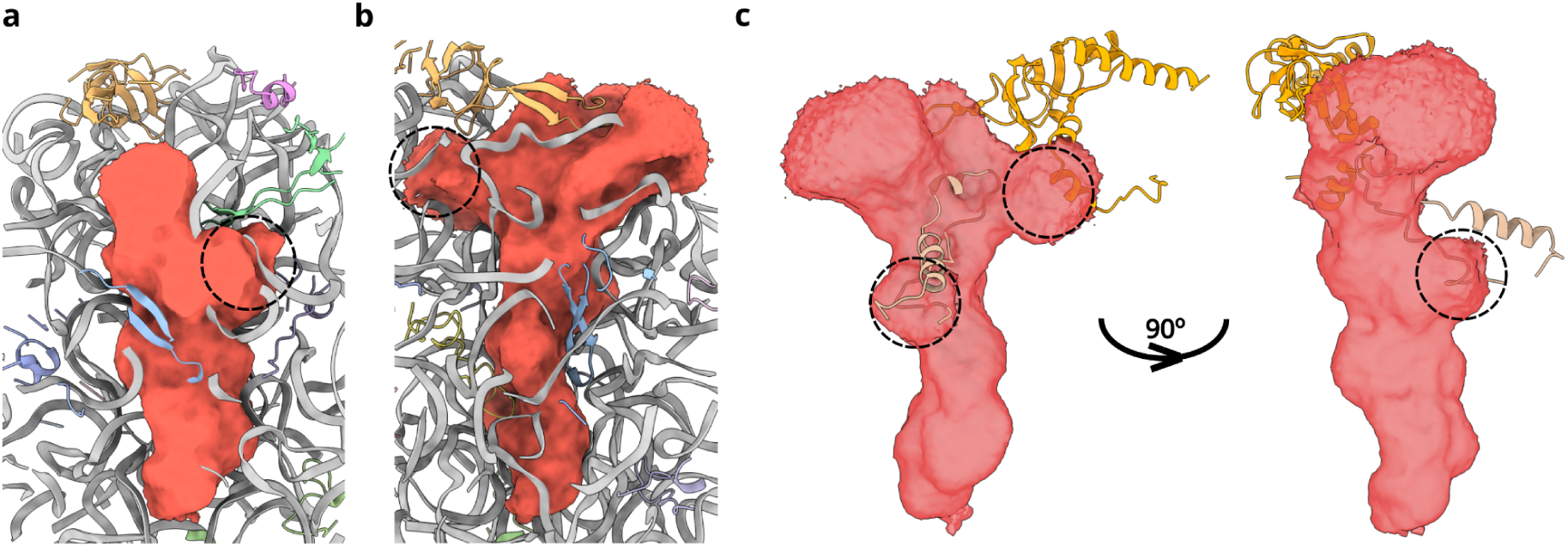
Characterisation of bacterial side tunnels and their evolutionary occlusion. **(a)** Location of the first bacteria-specific side tunnel (black dotted circle) relative to the functional tunnel volume (red density) at a nascent chain length of L=30. **(b)** Location of the second side tunnel (black dotted circle) at L=40. Ribosomal proteins are colored: uL4 (yellow), uL22 (blue), uL23 (green), uL24 (orange), and uL29 (violet). **(c)** Superposition of eukaryotic *(H. sapiens*) proteins eL39 (light brown) and uL24 (orange) onto the *E. coli* functional tunnel volume (L=40, red density), demonstrating the steric obstruction of the bacterial side tunnel loci by eukaryotic-specific elements.

To quantify the resulting deviation from a simple tube-like geometry, we calculated 2D asphericity of the tunnel cross-sections (Fig. 2f). Asphericity values near 0 indicate circular shapes, while values near 1 indicate highly elongated or irregular cross-sections. Across bacterial tunnels, asphericity values ranged from 0 to 0.55, confirming that a simple cylindrical tube model is not an accurate representation of the ribosomal exit tunnel geometry. The regions of highest irregularity coincide with the first constriction site and the two newly identified side tunnels. Conversely, the most circular regions are found close to the PTC, within the vestibule, and at ∼42 Å from the PTC (Fig. 2f), formed by uL22 and uL4 loops as well as rRNA.

Taken together, these results demonstrate that bacterial ribosome tunnels possess both highly conserved features, such as constriction sites, and distinct bacteria-specific secondary tunnels that are accessible only at specific stages of protein synthesis. These features influence the progression of nascent chains differently depending on chain length and position within the tunnel. The size, shape, and accessibility of these regions are likely influenced not only by sequence variation in ribosomal proteins and rRNA but also by post-translational and post-transcriptional modifications, as well as local dynamics. These findings reveal a nuanced and temporally regulated landscape that modulates the movement of nascent protein chains through the bacterial ribosome tunnel.

### Archaeal ribosome tunnels exhibit distinct structural adaptations

We next analysed the exit tunnels of five archaeal ribosomes representing two major phyla: Euryarchaeota and Thermoproteota (Fig. S10 and S11). As in bacteria, the functional volume of the tunnel changes as the nascent chain lengthens (Fig. 2b and S11), although the transitions appear less pronounced in archaea. The unique structural features of archaeal tunnels become especially apparent when compared to their bacterial counterparts (Fig. 2d).

Near the PTC, archaeal tunnels closely resemble bacterial ones, reflecting strong evolutionary conservation at both the sequence and structural levels (Fig. 2d). However, the first constriction site (∼30 Å from the PTC) is notably wider in archaea (Fig. 2d). This difference likely arises from the absence of a conserved arginine (equivalent to *E. coli* Arg61) that protrudes into the tunnel in bacteria but is missing in archaea (Fig. S12a). Moving further downstream, the tunnel architecture diverges significantly due to the incorporation of the archaeal/eukaryotic-specific protein eL39 and the presence of a distinct second loop in uL4 (absent in bacteria). These elements effectively remodel the tunnel wall, shifting the second constriction site ∼5 Å downstream (to ∼50 Å from the PTC) compared to bacteria (Fig. 2b, Fig. S13c).

Crucially, the structural addition of eL39 acts as an evolutionary “plug” that seals both lateral cavities observed in bacteria (Fig. 3c). The region corresponding to the bacterial first side tunnel is mainly occluded by eL39 (Fig. 3c and Fig. S12b), which packs against uL22 and rRNA, creating a narrower constriction in archaea than in bacteria at this depth (Fig. 2d). Similarly, the second side tunnel (60–70 Å) is effectively sealed by eL39 with the help of archaeal/eukaryotic-specific N-terminal extension to the uL24 (Fig. 3c), resulting in a third, highly pronounced constriction (Fig. 2b and S12c). Notably, the positioning of eL39 in this region directly intersects with the H6/H7 nascent chain pathway recently characterised in other systems (19, 41), suggesting that this specific pathway may be structurally occluded in the archeal ribosomes.

Despite this general trend toward a “sealed” tunnel, we observe two archaeal species that deviate from the general tunnel architecture. In *T. kodakarensis,* an unusual 23S rRNA conformation (nucleotides 2170-2173) blocks a significant portion of the tunnel near the PTC, obstructing the nascent chain’s path (Fig. S13a). This unique feature is also present in other available structures of this ribosome (PDB IDs: 6skf and 6skg (45)) but is distinct from the typical archaeal geometry near the PTC, as visible in our RMSD analysis (Fig. S1a). Conversely, *S. acidocaldarius* retains a side tunnel at a position similar to the bacterial ribosome (L=40) (Fig. S10), formed by a different conformation of rRNA and eL39 residues (Fig. S13b). Given the limited number of high-resolution archaeal ribosome structures (five species from two phyla), it remains an open question whether these features represent broad lineage-specific diversity or species-specific adaptations.

Finally, the tunnel asphericity in archaea also changes along the tunnel axis (Fig. 2f). The region around the PTC is very similar to the bacterial one, highlighting the structural and sequence conservation of this core region. The main differences correspond to the regions of bacterial-specific cavities, which are more spherical in archaea (at NC lengths of 30 and 40 amino acids). Notably, the region near the second constriction site exhibits significant differences in asphericity, despite less pronounced variations in cross-sectional area (Fig. 2d, e). This difference arises from the insertion of the second uL4 loop, which alters the shape - but not necessarily the overall cross-sectional area (Fig. S13c).

### Structural features of eukaryotic ribosome tunnels and the divergence of Microsporidia

Eukaryotic ribosomes comprise the largest group in our analysis, including 33 unique structures from 11 phyla (Ascomycota, Nematoda, Arthropoda, Chordata, Microsporidia, Euglenozoa, Fornicata, Streptophyta, Apicomplexa, Ciliophora and Parabasalia). While the majority of these structures share a highly conserved architecture, our initial analysis revealed that ribosome tunnels from Microsporidia (*E. cuniculi*, *P. locustae*, *S. lophii*, and *V. necatrix*) exhibit distinct structural features (Fig. S14, Fig. S15), consistent with recent observations (40). Consequently, we analysed these parasitic species separately from the canonical eukaryotic consensus.

Focusing first on the canonical eukaryotic ribosomes, we generated an averaged tunnel cross-sectional area profile, which showed the least variation across the three domains of life (Fig. 2c, Fig. S16). Notably, the overall architecture of the eukaryotic tunnel closely mirrors that of archaea, preserving three constriction sites consistently located at ∼30 Å, ∼50 Å and ∼67 Å from the PTC (Fig. 2c). These narrow regions are formed by the same structural elements found in archaea, most notably the eL39 protein, which occludes the regions corresponding to the two bacterial side tunnels (Fig. 3 and Fig. S17). This high degree of structural similarity between eukaryotic and archaeal exit tunnels, evident in both cross-sectional area and asphericity profiles (Fig. 2f), highlights their common evolutionary roots and confirms that the “sealed” tunnel architecture is a shared evolutionary trait.

In contrast, Microsporidia ribosomes exhibit markedly different tunnel architectures, reflecting the extensive ribosomal reduction typical of their obligate parasitic lifestyle. The extent of ribosome reduction varies across species, ranging from the relatively canonical tunnel of *P. locustae* to the heavily reduced structure in *V. necatrix* (Fig. S18). These reductions primarily affect the lower portion of the tunnel, beginning ∼60 Å from the PTC, which becomes significantly more open and solvent-exposed compared to other eukaryotes (Fig. S15 and S19). This aligns with recent geometric clustering that grouped specific parasitic protists with prokaryotes rather than canonical eukaryotes (40). While this increased architectural opening may facilitate nascent chain flexibility and earlier folding (19, 20), it likely also exposes the emerging polypeptide to premature interaction with cytosolic factors.

Taken together, our analysis reveals that cytoplasmic ribosome exit tunnels show a high degree of structural conservation within domains but can diverge fundamentally between lineages. This striking structural similarity between archaeal and eukaryotic tunnels, evident in cross-sectional area, asphericity profiles, and the specific “sealing” role of eL39, underscores their shared evolutionary history. In contrast, bacterial tunnels exhibit greater structural diversity, particularly in the variable opening of the two lineage-specific side tunnels, contributing to a broader architectural plasticity. This suggests an evolutionary trajectory in which the tunnel began as a variable, multi-branched landscape in bacteria but was refined into a more stringent, sealed conduit in the archaeal-eukaryotic lineage.

### The physicochemical landscape of the exit tunnel

Beyond geometric constraints, our computational approach also enabled us to explore the physicochemical environment experienced by the nascent chain as it traverses the ribosomal exit tunnel. We first mapped the rRNA-to-protein ratio along the tunnel path (L=60 NC simulations; Fig. 2e). While the tunnel is predominantly lined by rRNA, the central region, between the first and third constriction sites, shows a notable increase in protein content, particularly in archaea and eukaryotes. This enrichment arises from the presence of extended loops of uL4, uL22, and uL23 (in bacteria), as well as the eukaryotic- and archaeal-specific second uL4 loop and eL39 protein. Consequently, the tunnel wall in archaea and eukaryotes presents a significantly more protein-rich surface than its bacterial counterpart.

To further dissect the tunnel composition, we calculated the number of rRNA nucleotides and specific amino acids (polar, hydrophobic, positively charged, negatively charged, and special (Gly, Pro, and Cys)) lining the main tunnel path (Fig. 4). Strikingly, the rRNA profiles are virtually identical across all three domains of life. This indicates that the RNA provides a universally conserved scaffold, while the physicochemical differences of the tunnel are driven almost exclusively by protein composition.

**Figure 4.**
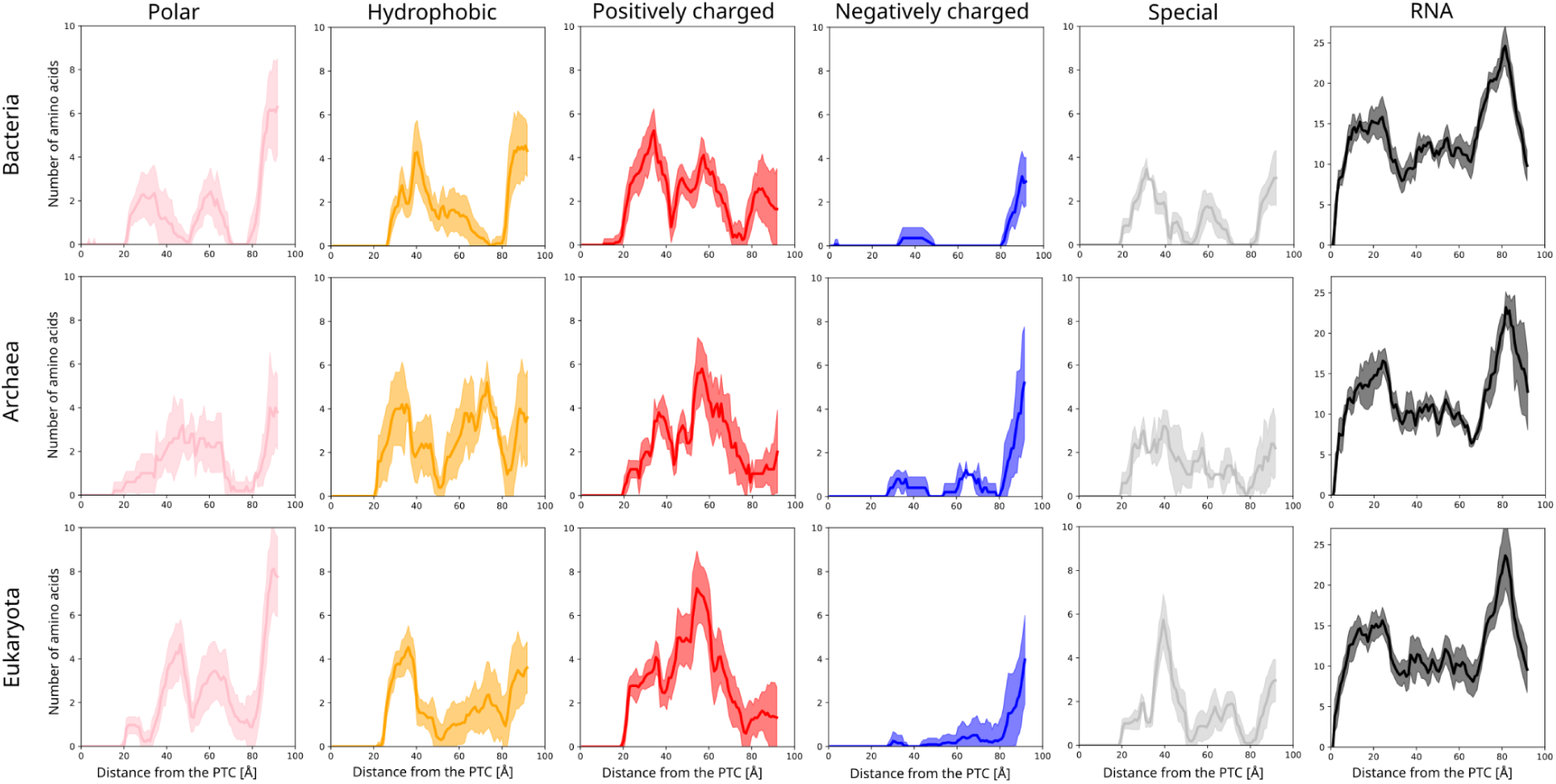
Divergent physicochemical landscape of the ribosomal exit tunnel. Comparison of the chemical environment lining the functional tunnel path across bacteria, archaea, and eukaryotes. The profiles display the contact frequency of rRNA nucleotides (top) and amino acid residues (bottom) as a function of distance from the PTC. Amino acids are grouped into five physicochemical categories: polar, hydrophobic, positively charged, negatively charged, and special (Gly, Pro, Cys).

Negatively charged residues are largely excluded from the tunnel interior, likely due to electrostatic repulsion from the dense, negatively charged rRNA backbone, and appear predominantly near the tunnel exit. Within the tunnel, the dominant residue types are polar, hydrophobic and positively charged (Fig. 4). Consistent with their structural homology, archaea and eukaryotic tunnels again display highly similar profiles, with the main difference observed in the region 60-70 Å from the PTC (the eL39 binding site). In the archaeal tunnel, this region is more hydrophobic and slightly less positively charged than in eukaryotes. In contrast, bacterial tunnels, which lack eL39, are generally less hydrophobic and positively charged in this region. An additional difference is observed at the first constriction site, where the bacterial tunnel shows an enrichment of polar residues, a feature consistent with the structural divergence noted earlier in this region.

## Discussion

In this study, we developed and applied a nascent chain-centric computational framework to define and analyse the cytoplasmic ribosome exit tunnel across the tree of life. Unlike previous geometry-based approaches (38, 40, 46), our method establishes a functional definition of the tunnel, based on the steric and entropic accessibility of the nascent polypeptide itself. This shift enables a natural delineation of tunnel boundaries and reveals transiently accessible features, most notably the bacteria-specific side tunnels that have not been previously characterised.

We demonstrate that the tunnel has a highly complex and irregular geometry, characterised by a non-linear axis and variable, non-circular cross-sections (high asphericity) (Fig. 2). This topological complexity precluded the use of simple radial measurements to accurately characterise tunnel dimensions. Instead, we propose functional cross-sectional area as the biologically relevant metric, which provides a more accurate proxy for the space accessible to the nascent chain. Consequently, our area-derived values are generally larger than those inferred from the commonly cited 10–20 Å radius range, as they account for irregular cavities, pockets and side-tunnels along the path.

Our functional mapping provides a mechanistic explanation for the geometric divergence observed between domains. While previous studies described the eukaryotic tunnel as “narrower” due to a second constriction (38), our analysis reveals that this narrowing results also from a specific evolutionary strategy: the “plugging” of ancestral side cavities. We show that the lateral branches accessible in bacteria are effectively sealed in archaea and eukaryotes by the incorporation of eL39 and an N-terminus extension of the protein uL24 (Fig. 3). This suggests an evolutionary trajectory in which the tunnel transitioned from a multi-branched, porous landscape in bacteria to a strictly confined conduit in higher organisms. This architectural “tightening” aligns with the hypothesis that the eukaryotic tunnel acts as a selective filter to trap specific small proteins (39). Furthermore, our observation that parasitic Microsporidia “re-open” these distal regions supports the recent finding that tunnel geometry is plastic and can revert to a bacterial-like state under specific evolutionary pressures (33); however, our observation suggests that the nature of the widening of the tunnel is very different from that of bacteria.

The exit tunnel vestibule is a critical environment in which proteins can initiate folding into native or near-native compact structures (10, 11, 13). The ribosome actively modulates this process, not only through direct enthalpy-driven interactions with the growing nascent chain (47–51), which can destabilise the native state (19, 52), but also through entropic destabilisation of the disordered ensemble (53). By restricting the conformational space available to the unfolded chain, the tunnel lowers the conformational entropy of the disordered state, thereby stabilising compact folding intermediates and altering potential folding pathways (53–55). Consequently, the specific shape and dimensions of the exit tunnel directly define this entropic landscape, likely tuning the trajectory of cotranslational folding.

While structural variability may arise from different ribosome functional states (e.g., empty vs. occupied) or variations in experimental methods and conditions, our PTC RMSD analysis (Fig. S1) and low variance across cross-sectional profiles (Fig. 2d) suggest these effects are minimal. However, an open question remains regarding how ribosomal dynamics, especially side-chain motion, can modulate its accessibility to the NC in real time. For instance, we have already observed that small changes in side chain orientation can have a significant impact on the effective geometry experienced by the NC. Incorporating ribosomal flexibility into future simulations will allow us to capture these effects more directly, particularly as our strategy employs structure-based models, which are well-adapted to various MD scenarios and widely used in ribosome simulations (56, 57).

Finally, our current use of a polyalanine NC sequence, with N-terminal methionine (eukaryotes/archaea) or formylmethionine (bacteria), and steric-only interactions serves as a topological baseline. By comparing this reference map against system-specific maps that incorporate side-chain chemistry, electrostatics, and specific stalling sequences, we can begin to dissect the interplay between tunnel architecture and nascent chain identity. Combining this with ribosome dynamics will open a new window to study these systems. This also enables prediction of the effects of post-translational and post-transcriptional modifications, antibiotic binding, or regulatory nascent peptide sequences - a key step toward predictive models of ribosome-NC interactions.

## Methods

### Search strategy for unique ribosome structures

To comprehensively collect known ribosome structures from the Protein Data Bank, we used a dual-strategy approach that combined keyword-based and sequence-similarity searches. First, we performed a search using the term “ribosome” in the Structure Keywords field, refining the results to include only structures containing both protein and RNA components determined by either X-ray crystallography or single-particle cryo-electron microscopy. Second, as an orthogonal approach, we selected from the PDB all experimentally determined structures that contained at least one protein chain with ≥30% sequence identity to any known highly conserved uL2 protein. These two approaches yielded 1, 939 and 1, 438 PDB entries, respectively, resulting in a total of 2, 069 unique ribosome structures. From this dataset, we selected a single representative structure per organism, prioritising high resolution and the presence of nascent chain (translating ribosomes), while strictly excluding structures containing antibiotics or other factors bound in or near the ribosomal exit tunnel. For example, the available structures for *Cutibacterium acnes* were excluded due to the presence of tunnel-bound sarecycline (Table S5). Our final dataset comprised 55 unique ribosome structures, of which 17 were from Bacteria (Table S1), 33 from Eukaryotes (Table S2), and 5 from Archaea (Table S3).

### Definition of the reference ribosome tunnel

To build the reference model of the ribosome exit tunnel, we used our previously published *E. coli* ribosome structure containing the FLN5 filamin domain nascent chain (PDB ID: 7zp8 (19)). Initially, we selected all ribosomal residues within 30 Å of the available NC model using VMD (58) to define the preliminary tunnel boundaries. To capture the full extent of the ribosomal tunnel accessible to the nascent chain, we used all-atom structure-based molecular dynamics simulations. The NC was modelled as a poly-alanine chain with an N-terminal formylmethionine (N-fMet) at five distinct lengths (L = 10, 20, 30, 40, and 60 amino acids). These lengths were chosen to represent various stages of translation; the L=40 chain was sufficient to span the entire length of the tunnel, whereas the L=60 chain allowed sampling of the ribosome surface surrounding the exit tunnel. All-atom structure-based models were generated using SMOG2 (template: SBM_AA-amber-bonds) (59–61) for these five RNCs. For each system, 100 ns simulations were conducted using GROMACS 2021 (see details below), with ribosomal atoms and the C-terminal residue of the NC held fixed. We mapped the resulting MD trajectories onto the full *E. coli* ribosome structure (PDB ID: 7zp8 (19)) and, using a 6 Å cut-off, identified additional tunnel residues that were missed in the initial selection. This approach provided a more comprehensive definition of the reference ribosome tunnel.

### Structural alignment and selection of the ribosome exit tunnels

The reference ribosome exit tunnel from *E. coli* was used to guide the selection of the exit tunnels from the remaining 54 ribosome structures. Given the complexity of aligning whole ribosome structures across diverse organisms, we leveraged available cryo-EM maps to facilitate the process. We initially performed a rigid-body fit of each target ribosome structure into the cryo-EM density map of the reference *E. coli* ribosome (EMDB 14850) using ChimeraX. Following this global fit, we performed a localised refinement using a simulated density map of the reference *E. coli* ribosome tunnel, generated in ChimeraX at 6 Å resolution. This step optimised the alignment specifically within the exit tunnel region.

To validate this density-based alignment, we calculated the RMSD of the conserved PTC residues (as defined in (62)) between the reference *E. coli* structure and each aligned ribosome (Supplementary Figure 1A). Once aligned, tunnel-lining residues were identified in each structure using VMD by selecting atoms located within the volume of the reference *E. coli* tunnel density. All selections were visually validated in ChimeraX; in cases of significant structural divergence, the selection was manually extended based on visual inspection in VMD.

### Modelling of the missing residues of the exit tunnels

The majority of the generated ribosome tunnel structures (44/55) were complete; however, in a few cases, modelling of missing regions was required (Table S4). For ribosomes from *D. radiodurans*, *F. johnsoniae*, and *S. aureus*, a single nucleotide (U2585, U2583, and U2612, respectively) was removed as it overlapped with the C-terminal residue of the nascent chain. This minor modification did not affect the functional shape of the tunnel. The ribosome structures from *F. johnsoniae*, *E. cuniculi*, *G. lamblia*, *N. crassa*, *P. falciparum*, *S. lophii*, *T. gondii*, *V. necatrix*, and *H. marismortui* contained gaps in various protein or rRNA regions within the tunnel, which were modelled using AlphaFold 2 (63) and AlphaFold 3 (64). Additionally, certain structures required the removal of obstructing factors prior to simulation: the dormancy factor MDF2 was removed from the *V. necatrix* ribosome, the protein Dap1 was removed from the *X. laevis* tunnel, and a nascent chain density was removed from the *R. norvegicus* structure.

### Building starting models for the MD simulations

For each ribosome exit tunnel model, we generated five RNC starting structures with poly-alanine nascent chains of distinct lengths: 10, 20, 30, 40, and 60 amino acids. The N-terminal residue was modelled based on the ribosome’s biological origin: methionine for eukaryotic and archaeal ribosomes and formylmethionine for bacterial, consistent with the translation initiation in each organism. A proline residue was placed at the C-terminus, following the initial NC model from the *E. coli* ribosome structure (PDB ID: 7zp8) (19), and was held fixed during the simulation. Certain ribosome structures required additional modifications to ensure compatibility with the CHARMM force fields used in the structure-based model simulations. Specifically, if parameters for a post-transcriptional modification were unavailable in CHARMM, the residue was mutated back to its unmodified state.

### MD simulations of the ribosome nascent chain complexes

Each RNC was simulated three times (replicas) for 100 ns in GROMACS 2021 (65) using an all-atom structure-based model generated via SMOG2 (template: SBM_AA-amber-bonds) (59–61). The starting NC conformation for the first replica was derived from the reference nascent chain (PDB ID: 7zp8), with minor manual adjustments in Coot to resolve steric clashes as needed. To enhance sampling efficiency, starting NC conformations for the two subsequent replicas were selected from the initial trajectory, based on backbone RMSD; we specifically chose frames that were structurally most distinct from each other and from the initial starting structure. This approach ensured that each simulation was initiated from a very different point in conformational space, facilitating robust sampling of the exit tunnel volume. Throughout the simulations, the ribosomal atoms were held fixed, and the interactions between the NC and the ribosome were treated as purely steric.

### Properties of cytoplasmic ribosome exit tunnels

To systematically compare the structural and physicochemical features of ribosome exit tunnels across bacteria, archaea and eukaryotes, we defined a single reference tunnel path that captures the common exit pathway shared by these ribosomes. We selected the tunnel centerline from the microsporidian *Encephalitozoon cuniculi,* as its geometry is relatively linear and free of major twists or bends within and beyond the vestibule (Fig. S5). This linearity minimises distortions when calculating the perpendicular cross-sectional area along the tunnel, enabling more accurate dimensional comparisons. In addition to cross-sectional area, which serves as our proxy for the tunnel geometry, we examined how key molecular properties change along this reference path. Specifically, we mapped the spatial distribution of polar, hydrophobic, positively and negatively charged residues and chemically distinct amino acids (Gly, Cys, and Pro). We also calculated the local protein-to-RNA composition ratio, providing insight into the changing molecular environment encountered by the nascent chain during translation.

### Trajectory analysis

In our approach, we used the NC dynamics to define the functional shape of the ribosomal tunnel. Each NC trajectory was aligned to the starting conformation using the C-terminal proline residue as a reference, enabling direct structural comparison across all systems. Subsequently, each replica trajectory was converted into a volumetric occupancy map using a defined 3D grid (voxel size = 1Å), from which an average occupancy map was calculated. This average map effectively serves as a “plaster cast” of the ribosomal tunnel, delineating all the regions accessible for the NC inside the ribosome. We divided this volume into “edge” voxels that follow the contours of the tunnel walls, and “interior” voxels, based on their location within the map. The edge voxels were used to identify the ribosomal residues lining the tunnel, which were then classified into polar, hydrophobic, positively charged, negatively charged, special residues (Gly, Pro and Cys) and RNA. Additionally, from each trajectory, we determined the path of maximum occupancy, defined by the voxels within the maximum occupancy for each residue (Cɑ atoms). We then applied cubic B-spline interpolation to this pathway to refine the main path inside the exit tunnel and calculated the average pathway for each ribosome across the three replicas. Using this averaged tunnel path, we defined the tunnel cross-sections as a disc combining both edge and interior voxels, perpendicular to each point along the tunnel axis.

### Data Availability

All raw and processed data resulting from the molecular dynamics simulations, including MD trajectories and the 55 species-specific occupancy maps and calculated cross-sectional area profiles, have been deposited in Zenodo and can be accessed at 10.5281/zenodo.18079792.

## Acknowledgements

This research is part of the project No. 2022/47/P/NZ1/03127 co-funded by the National Science Centre and the European Union’s Horizon 2020 research and innovation programme under the Marie Skłodowska-Curie grant agreement no. 945339. For the purpose of Open Access, the author has applied a CC-BY public copyright licence to any Author Accepted Manuscript (AAM) version arising from this submission. We gratefully acknowledge the Polish high-performance computing infrastructure PLGrid (HPC Centre: ACK Cyfronet AGH) for providing computer facilities and support within the computational grant no. PLG/2024/017461.

