## Supplementary Figures for "Evolution of the ribosomal exit tunnel through the eyes of the nascent chain"

A)

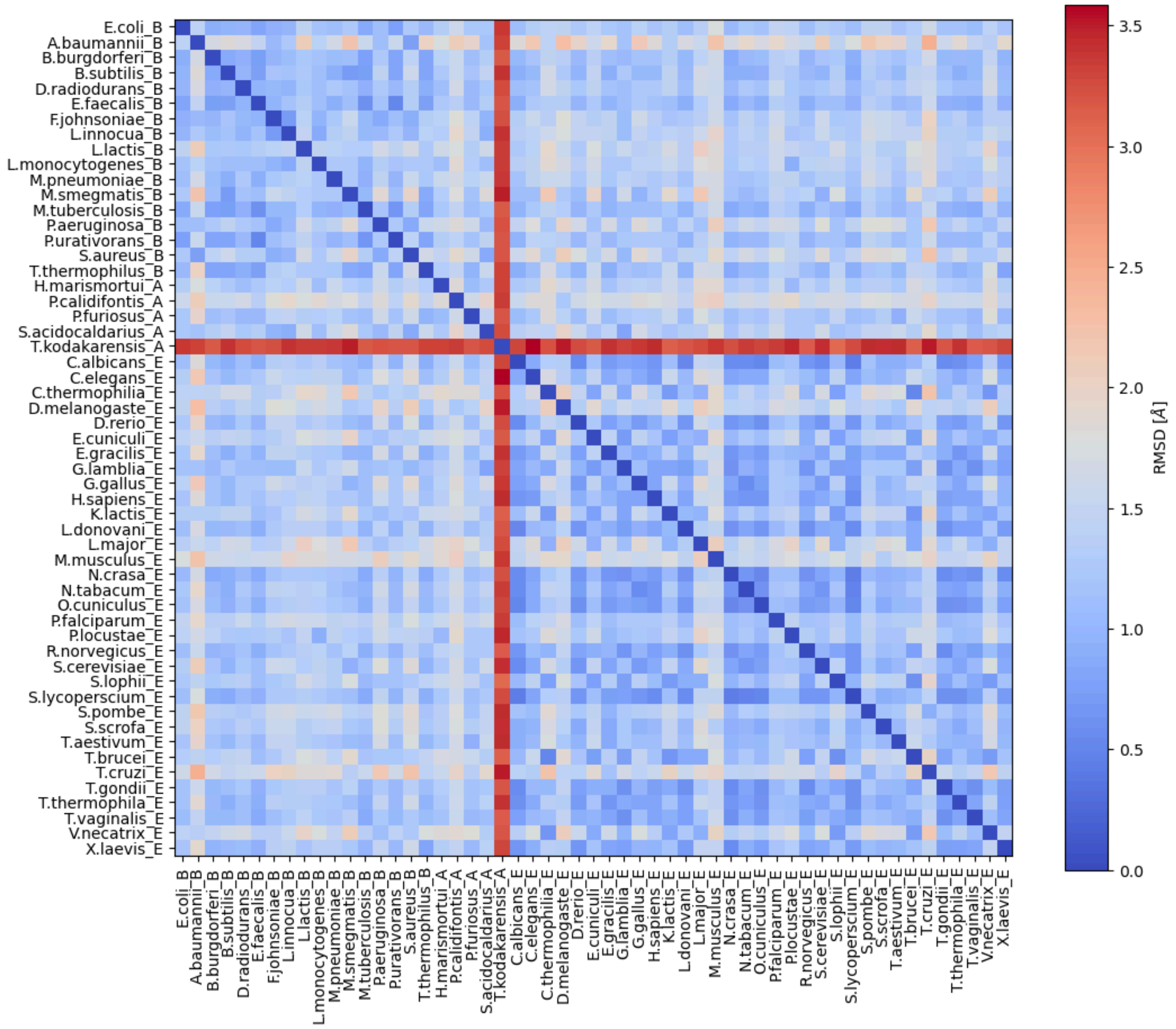

B)

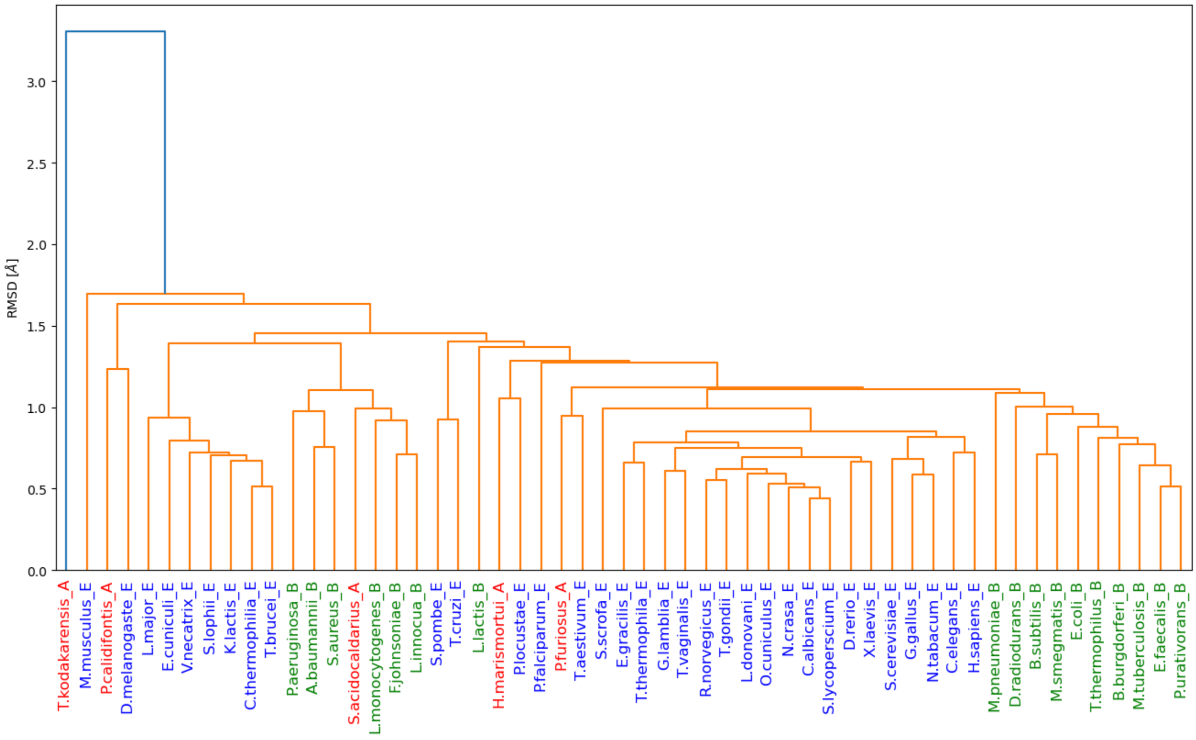
RMSD [Å]
0.0
0.5
1.0
1.5
2.0
2.5
3.0
T.kodakarensis_A
M.musculus_E
P.calidifontis_A
D.melanogaste_E
L.major_E
E.cuniculi_E
V.necatrix_E
S.lophii_E
K.lactis_E
C.thermophilia_E
T.brucei_E
P.aeruginosa_B
A.baumannii_B
S.aureus_B
S.acidocaldarius_A
L.monocytogenes_B
F.johnsoniae_B
L.innocua_B
S.pombe_E
T.cruzi_E
L.lactis_B
H.marismortui_A
P.locustae_E
P.falciparum_E
P.furiosus_A
T.aestivum_E
S.scrofa_E
E.gracilis_E
T.thermophila_E
G.lamblia_E
T.vaginalis_E
R.norvegicus_E
T.gondii_E
L.donovani_E
O.cuniculus_E
N.crasa_E
C.albicans_E
S.lycoperscium_E
D.rerio_E
X.laevis_E
S.cerevisiae_E
G.gallus_E
N.tabacum_E
C.elegans_E
H.sapiens_E
M.pneumoniae_B
D.radiodurans_B
B.subtilis_B
M.smegmatis_B
E.coli_B
T.thermophilus_B
B.burgdorferi_B
M.tuberculosis_B
E.faecalis_B
P.urativorans_B

C)

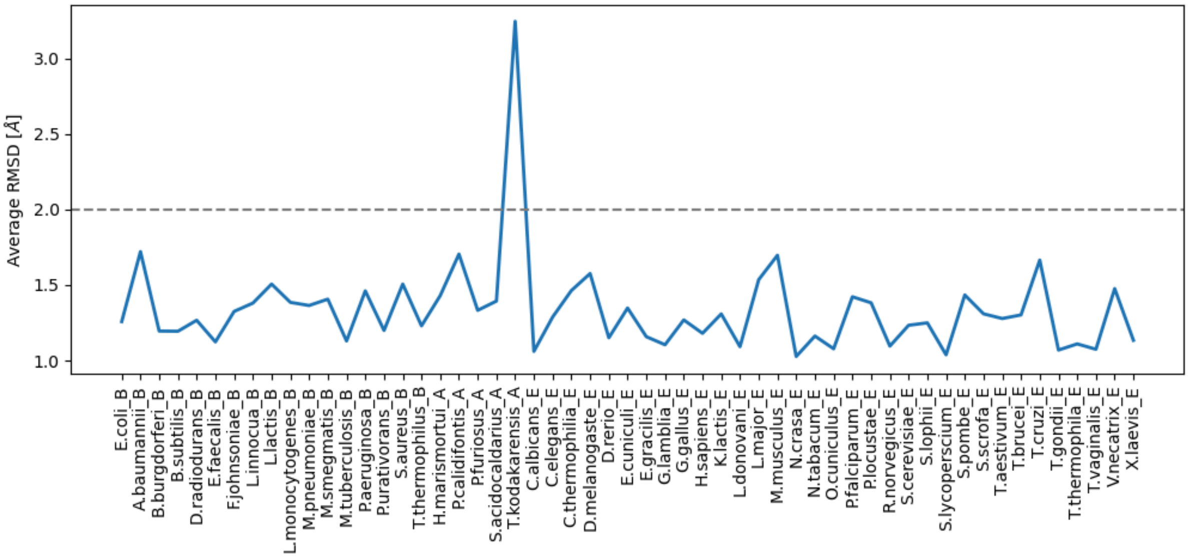
Average RMSD [Å]
1.0
1.5
2.0
2.5
3.0
E.coli_B
A.baumannii_B
B.burgdorferi_B
B.subtilis_B
D.radiodurans_B
E.faecalis_B
F.johnsoniae_B
L.innocua_B
L.lactis_B
L.monocytogenes_B
M.pneumoniae_B
M.smegmatis_B
M.tuberculosis_B
P.aeruginosa_B
P.urativorans_B
S.aureus_B
T.thermophilus_B
H.marismortui_A
P.calidifontis_A
P.furiosus_A
S.acidocaldarius_A
T.kodakarensis_A
C.albicans_E
C.elegans_E
C.thermophilia_E
D.melanogaste_E
D.rerio_E
E.cuniculi_E
E.gracilis_E
G.lamblia_E
G.gallus_E
H.sapiens_E
K.lactis_E
L.donovani_E
L.major_E
M.musculus_E
N.crasa_E
N.tabacum_E
O.cuniculus_E
P.falciparum_E
P.locustae_E
R.norvegicus_E
S.cerevisiae_E
S.lophii_E
S.lycoperscium_E
S.pombe_E
S.scrofa_E
T.aestivum_E
T.brucei_E
T.cruzi_E
T.gondii_E
T.thermophila_E
T.vaginalis_E
V.necatrix_E
X.laevis_E

**Figure S1. Structural conservation of the Peptidyl Transferase Centre (PTC) across phylogeny. (A)** Pairwise root-mean-square deviation (RMSD) matrix for the PTC region calculated across 55 ribosome structures representing Bacteria, Archaea, and Eukarya. **(B)** Hierarchical clustering dendrogram derived from the pairwise RMSD matrix using the average linkage method, which broadly recapitulates the separation between domains. **(C)** Average RMSD values for each of the 55 PTC structures, highlighting the high degree of structural conservation across the dataset with limited outliers (e.g., *T. kodakarensis*).

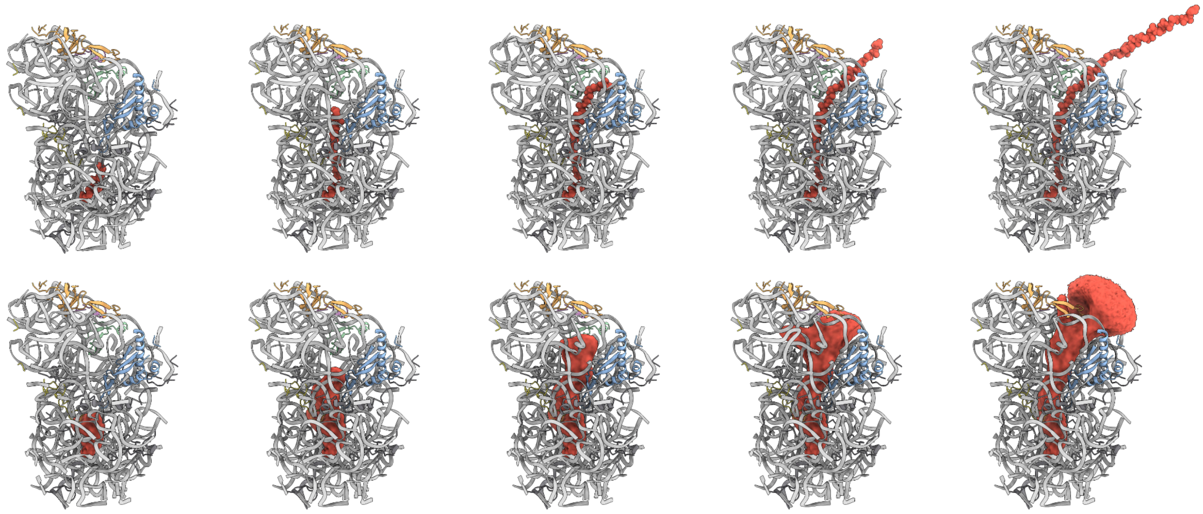

**Figure S2. Generation of functional tunnel maps using a nascent chain-centric approach. Top:** Initial structural models of RNC complexes used for simulations with polyalanine nascent chains of increasing lengths (10, 20, 30, 40, and 60 residues) were attached to the P-site tRNA to probe different biosynthetic time points. **Bottom:** Representative volumetric density maps derived from the all-atom MD simulations. These maps represent the spatial occupancy of the nascent chain, defining the functional volume of the tunnel accessible at a specific chain length.

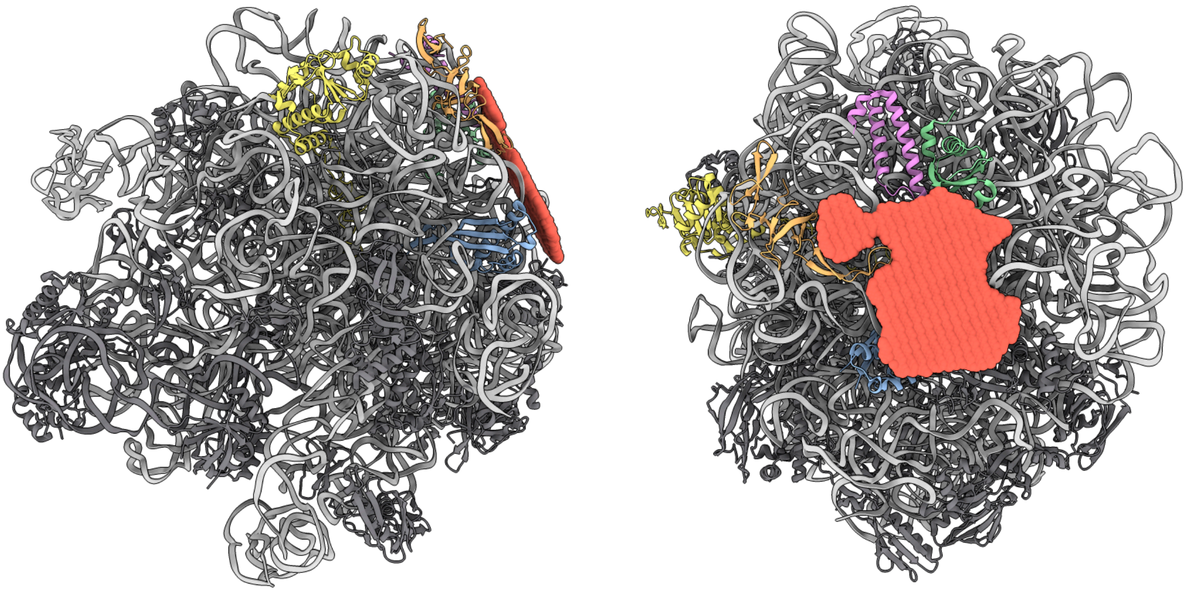

**Figure S3. Definition of the ribosomal tunnel exit boundary.** The lower boundary of the exit tunnel was defined by a density plane (red disc) aligned with the conserved loop of protein uL24. Key tunnel-lining proteins are colored for reference: uL4 (yellow), uL22 (blue), uL23 (green), uL24 (orange) and uL29 (violet).

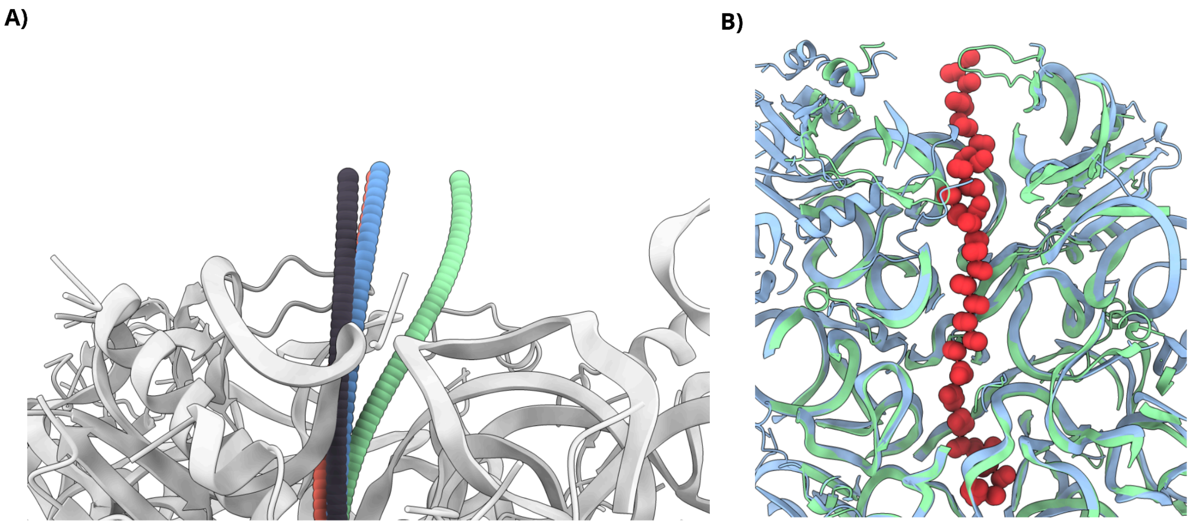


**Figure S4. Comparison of nascent chain trajectories and tunnel profiles. (A)** Superposition of nascent chain exit pathways for representative species: *E. coli* (green), *H. sapiens* (blue), *H. marismortui* (red), and *E. cuniculi* (black). **(B)** The cross-section of the ribosome exit tunnel along the main axis for Bacteria (green) and Eukarya (blue), overlaid with a representative nascent chain model (red).

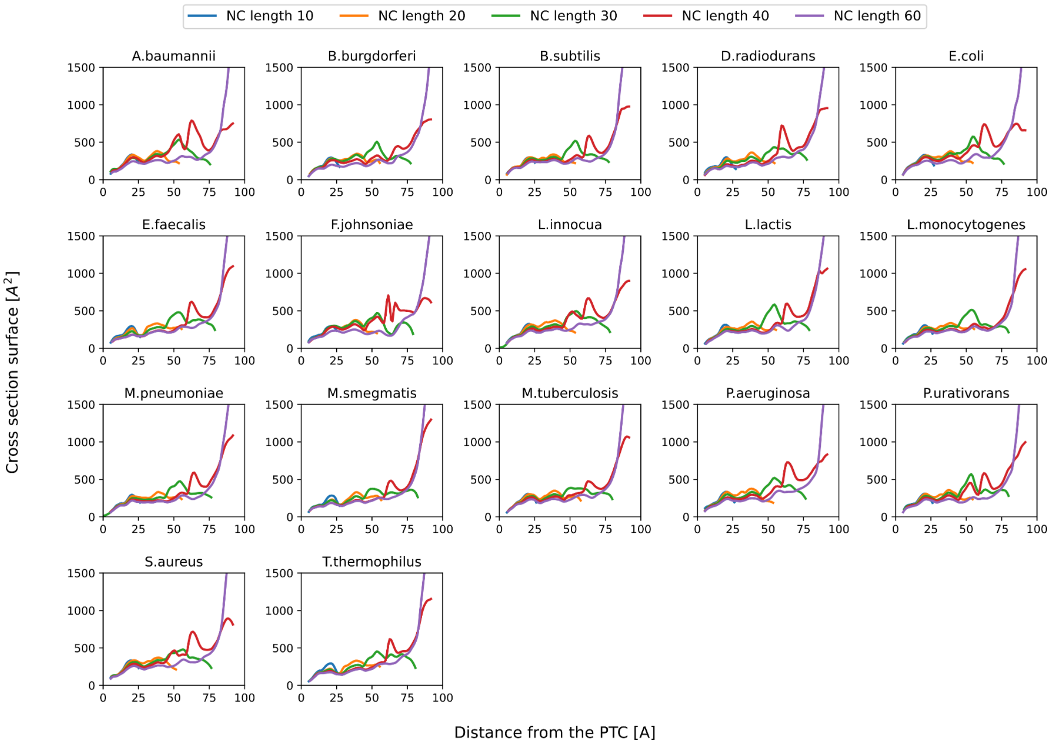


**Figure S5. Functional cross-sectional area profiles of bacterial ribosome exit tunnels.** Profiles were calculated for 17 bacterial structures using simulations with five increasing nascent chain lengths (L = 10, 20, 30, 40, and 60 residues), illustrating the length-dependent accessibility of the tunnel volume.

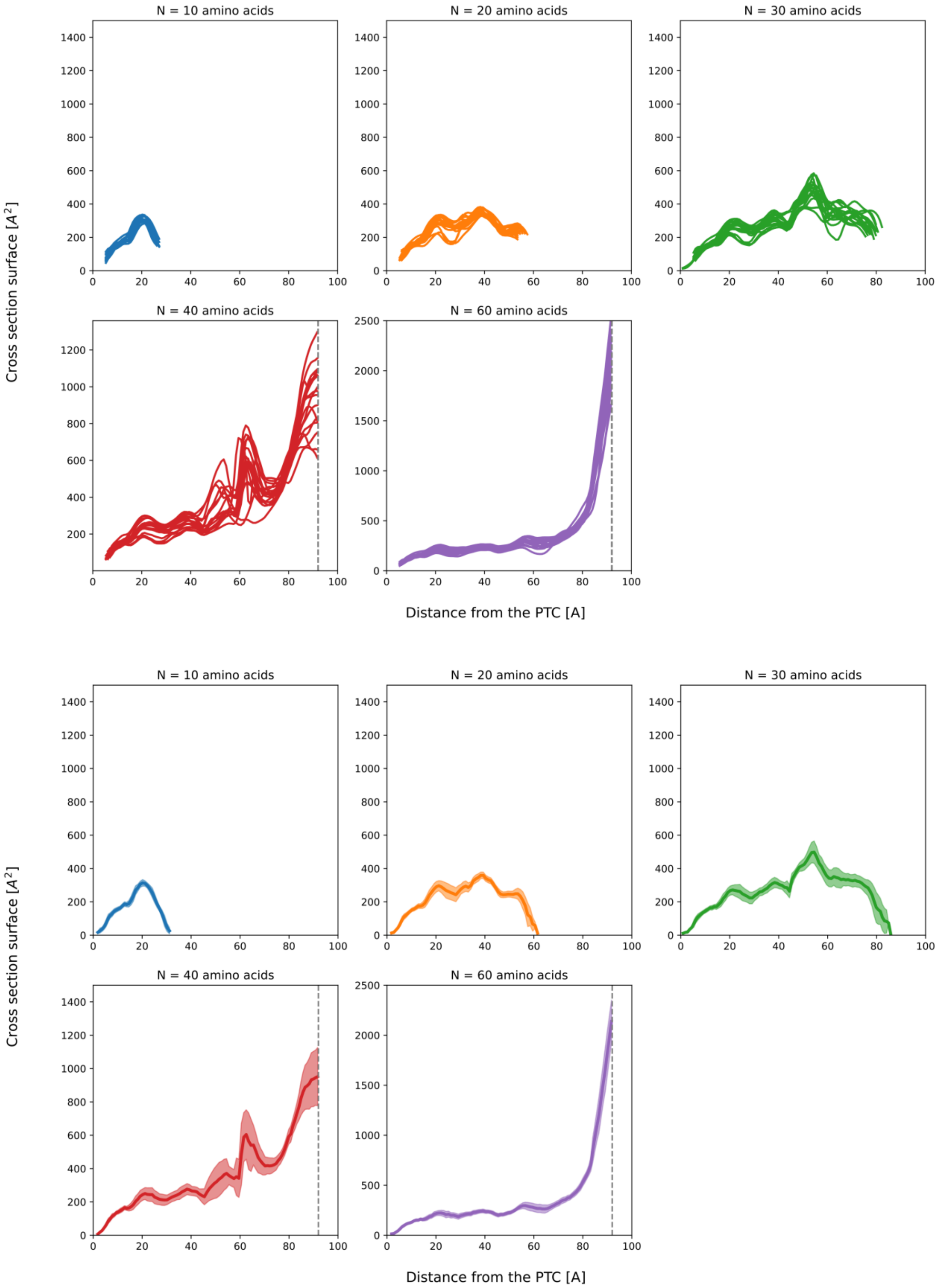


**Figure S6. Combined cross-sectional area profiles of bacterial ribosome tunnels. (Top)** Superimposed cross-sectional area profiles derived from all 17 analysed bacterial ribosomes across all nascent chain lengths, visualising the full range of structural variability. **(Bottom)** The average cross-sectional area profile for the bacterial dataset, with the shaded region representing the standard deviation (± SD). A vertical grey dotted line indicates the functional exit boundary of the tunnel.

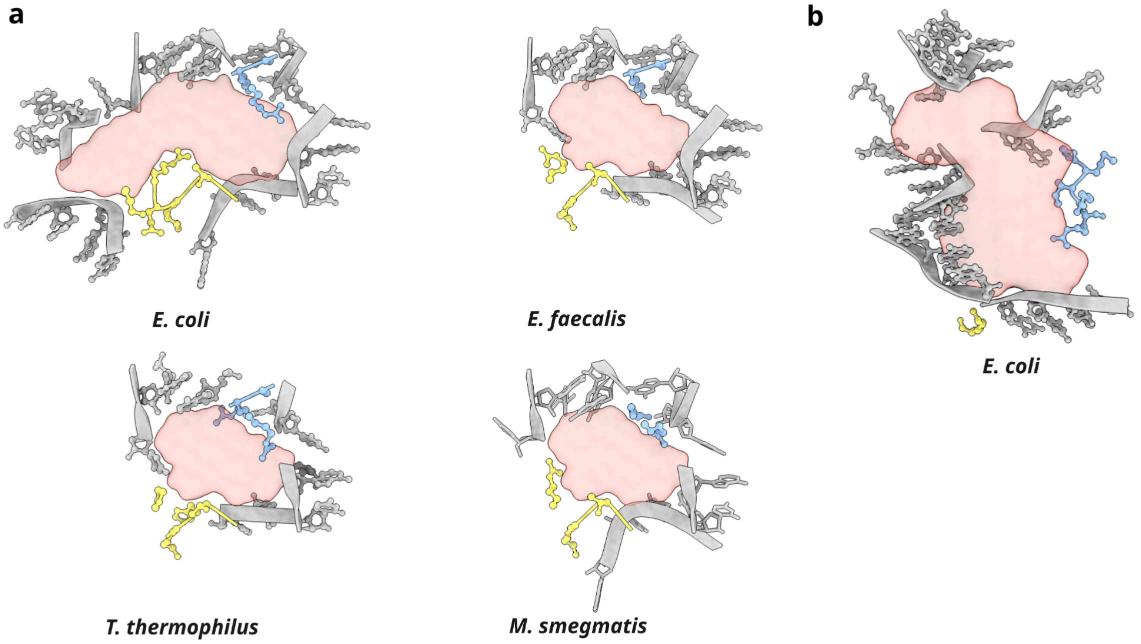


**Figure S7. Structural variations at bacterial constriction sites. (A)** Cross-sectional view of the first constriction site (~30 Å from the PTC) in *E. coli*, *E. faecalis*, *T. thermophilus*, and *M. smegmatis*, highlighting species-specific differences in tunnel geometry. **(B)** Structural detail of the second narrowing region in the *E. coli* ribosome. Ribosomal proteins are colored for reference: uL4 (yellow) and uL22 (blue).

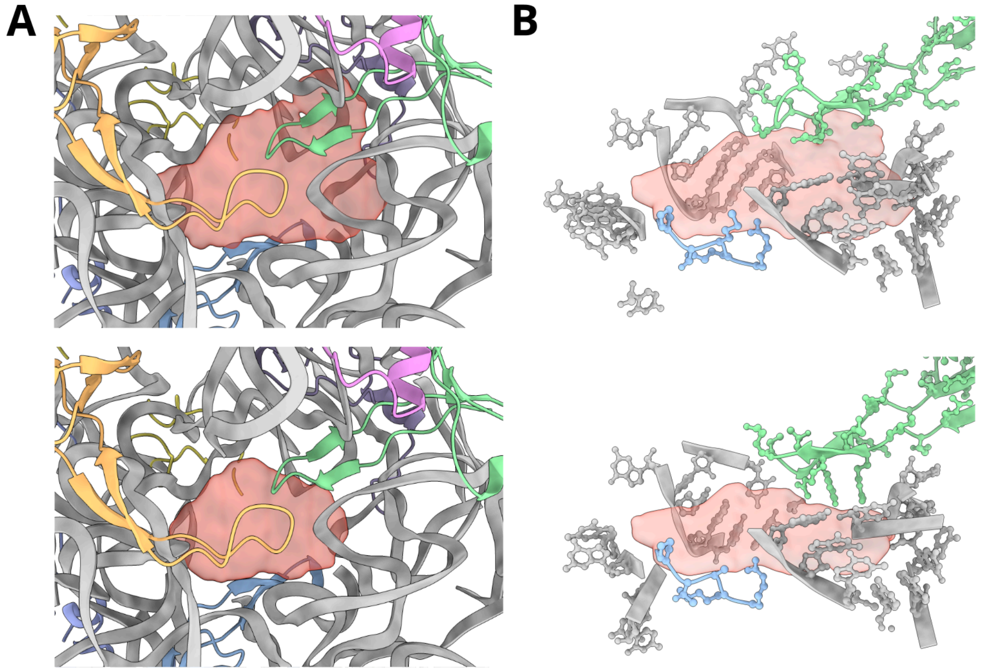


**Figure S8. Characterisation of the first bacterial side tunnel and its accessibility. (A)** Cross-sections of the first side tunnel region (~55 Å from the PTC) at nascent chain lengths of L=30 (top) and L=60 (bottom), illustrating the length-dependent gating of this cavity. **(B)** Species-specific structural variation in the first side tunnel (at L=30) between *E. coli* (top) and *M. tuberculosis* (bottom), highlighting local restriction in *M. tuberculosis*. Ribosomal proteins are colored: uL4 (yellow), uL22 (blue), uL23 (green), uL24 (orange), and uL29 (violet).

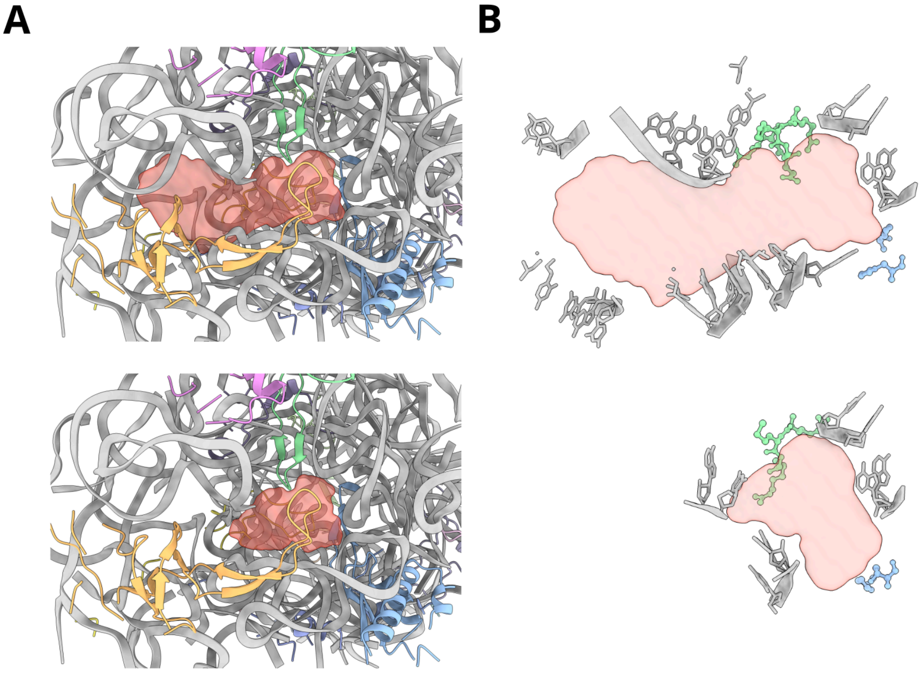


**Figure S9. Characterisation of the second bacterial side tunnel. (A)** Cross-sections of the second side tunnel region (~65 Å from the PTC) at nascent chain lengths of L=40 (top) and L=60 (bottom), illustrating changes in accessibility during elongation. **(B)** Structural occlusion of the second side tunnel in *L. monocytogenes*. Comparison of cross-sections at ~65 Å (L=40) between *E. coli* (top, open) and *L. monocytogenes* (bottom, closed). Ribosomal proteins are colored: uL4 (yellow), uL22 (blue), uL23 (green), uL24 (orange), and uL29 (violet).

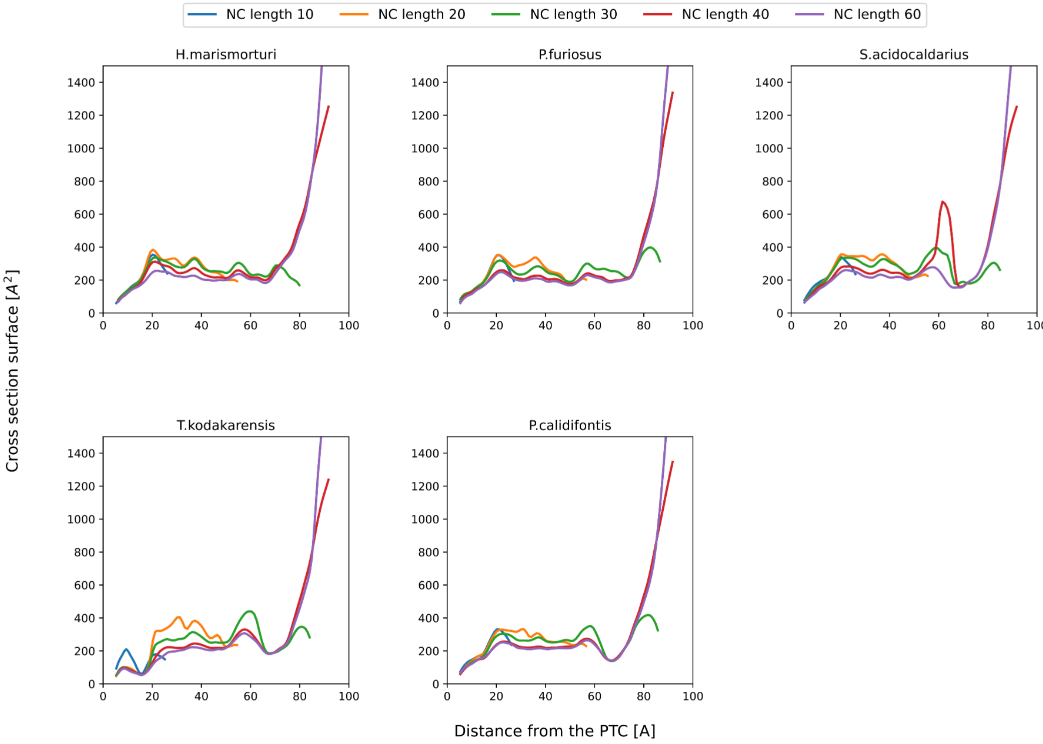

**Figure S10. Functional cross-sectional area profiles of archaeal ribosome exit tunnels.** Profiles were calculated for the five available archaeal structures using simulations with five increasing nascent chain lengths (L = 10, 20, 30, 40, and 60 residues), illustrating the length-dependent accessibility of the tunnel volume.

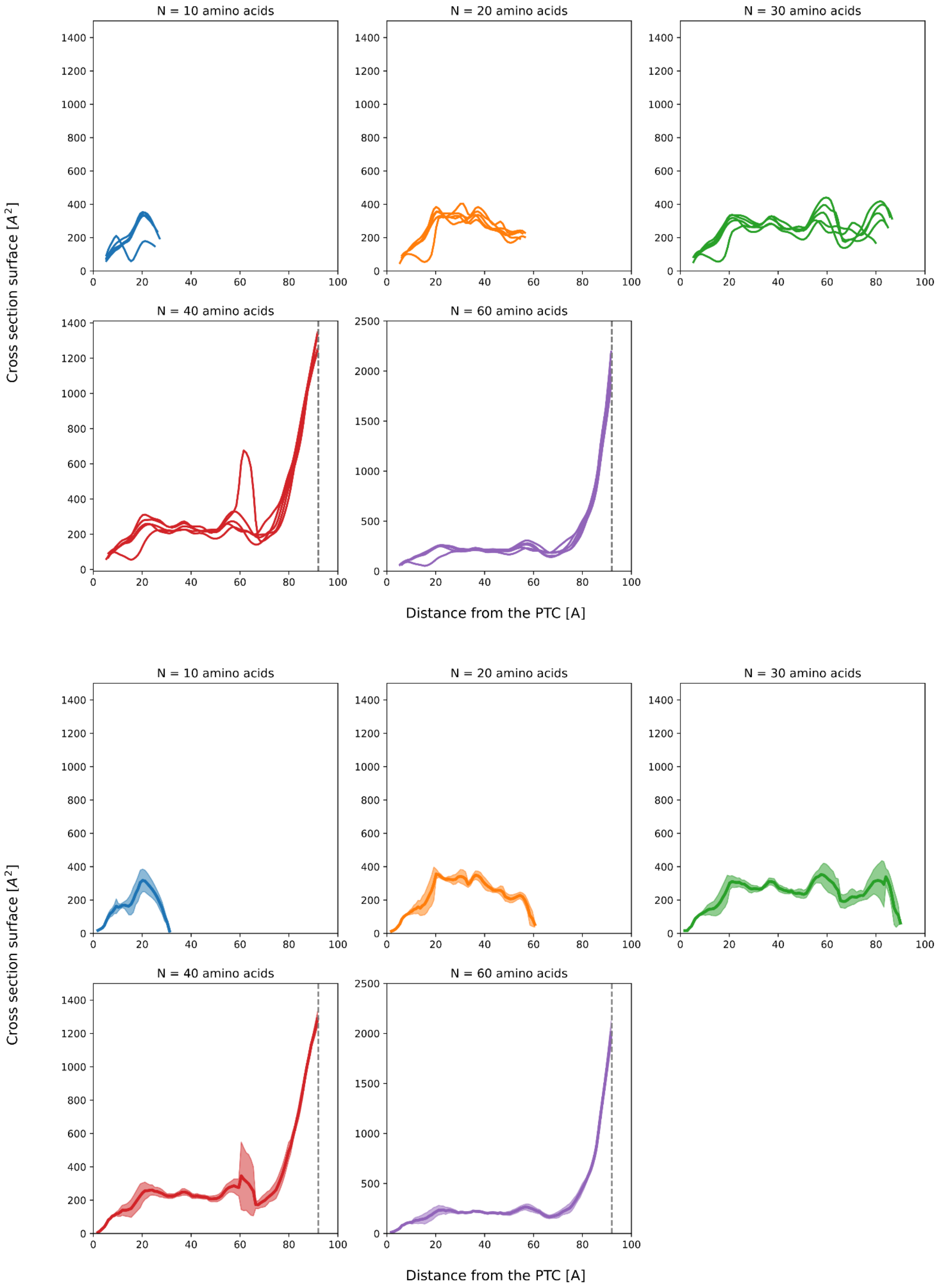

**Figure S11. Combined cross-sectional area profiles of archaeal ribosome tunnels. (Top)** Superimposed cross-sectional area profiles derived from all five analysed archaeal ribosomes across all nascent chain lengths, visualising the structural variability within the domain. **(Bottom)** The average cross-sectional area profile for the archaeal dataset, with the shaded region representing the standard deviation (± SD). A vertical grey dotted line indicates the functional exit boundary of the tunnel.

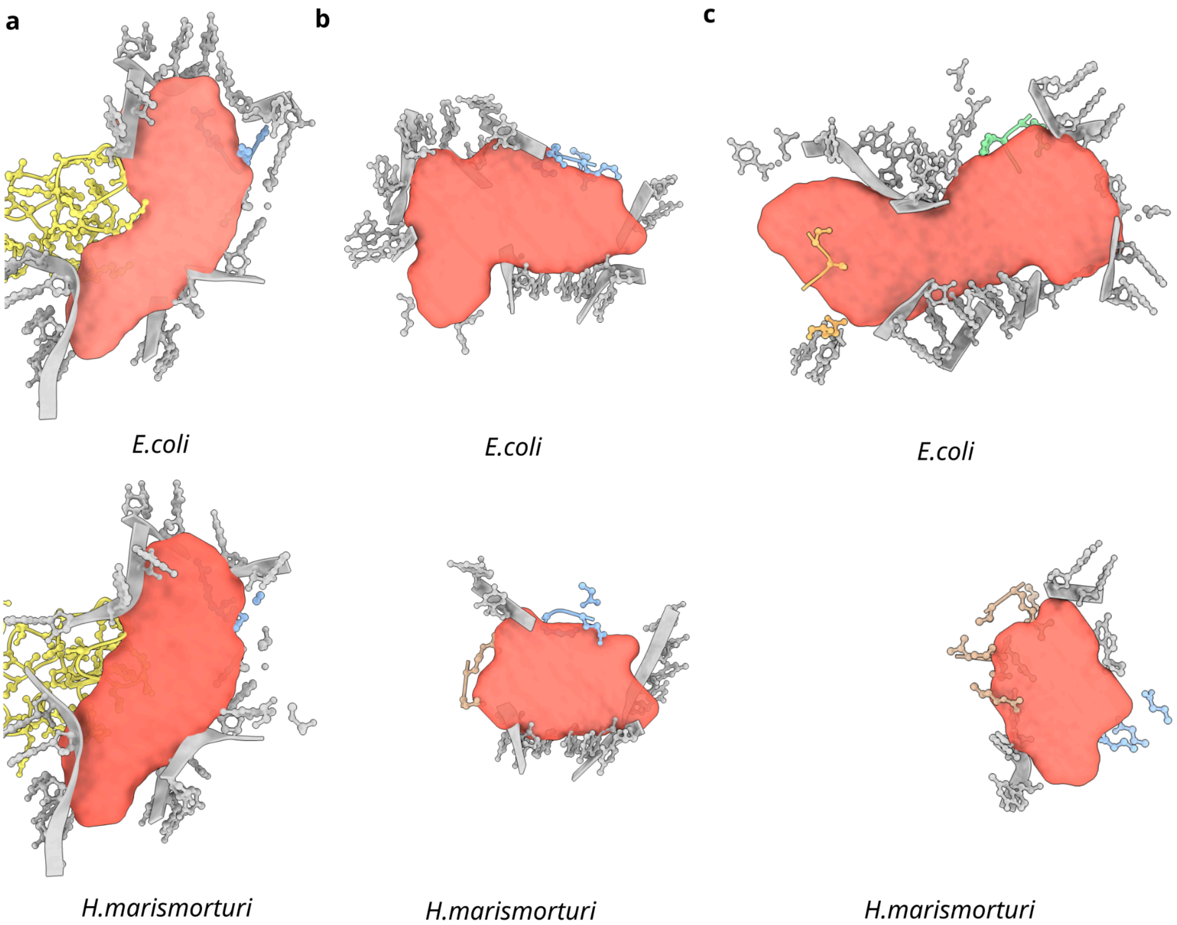


**Figure S12. Structural comparison of bacterial openings and archaeal constrictions. (A)** The first constriction site (~30 Å from PTC): Comparison between *E. coli* (top) and *H. marismortui* (bottom), showing the wider architecture in Archaea. **(B)** The second archaeal constriction region (~50 Å): *E. coli* (top) exhibits an open lateral cavity (first side tunnel), whereas *H. marismortui* (bottom) forms a constriction due to the presence of eL39 (light brown) and the uL4 loop (yellow). **(C)** The third archaeal constriction region (~67 Å): *E. coli* (top) shows the opening of the second side tunnel, while *H. marismortui* (bottom) is sealed by eL39. Ribosomal proteins are colored: uL4 (yellow), uL22 (blue), uL23 (green), uL24 (orange), and eL39 (light brown).

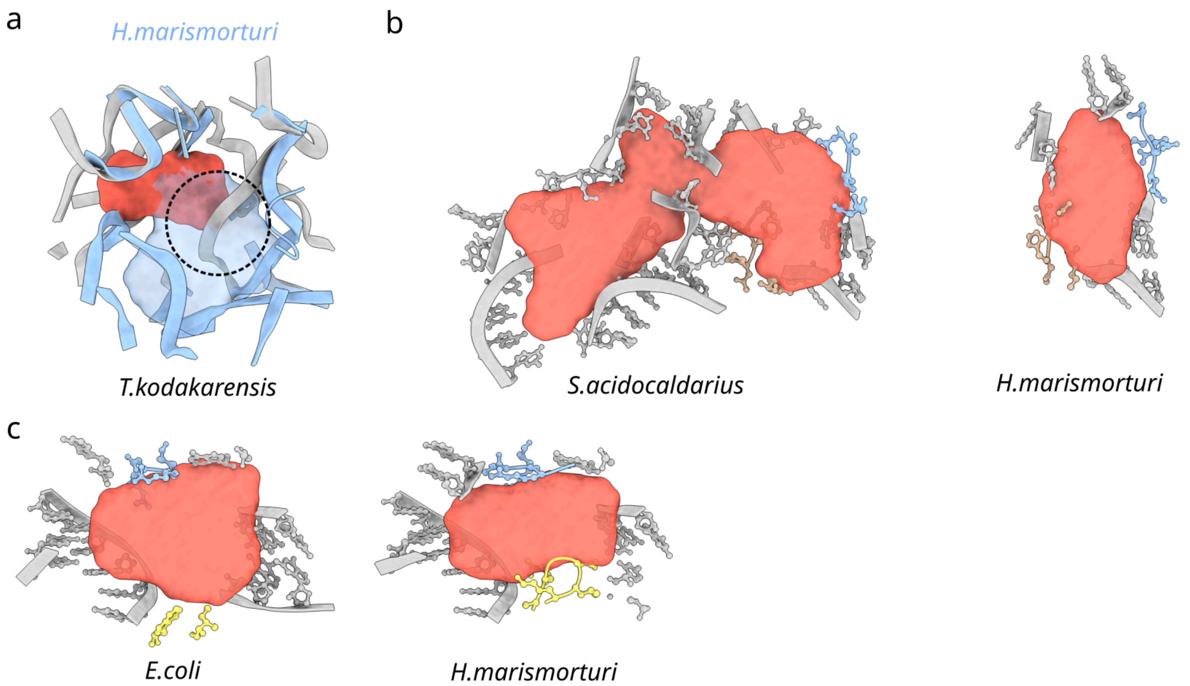


**Figure S13. Structural anomalies and adaptations in archaeal tunnels. (A)** Unique PTC obstruction in *T. kodakarensis*. Comparison of the PTC region in *H. marismortui* (left) and *T. kodakarensis* (right). The circled region highlights a distinct rRNA conformation in *T. kodakarensis* that sterically occludes the nascent chain path. **(B)** Atypical side tunnel retention in *S. acidocaldarius*. Cross-sections showing the "side tunnel" region in *S. acidocaldarius* (left, open) compared to the canonical "sealed" architecture of *H. marismortui* (right). **(C)** Modulation of tunnel geometry by the second uL4 loop. Visualisation of how the archaeal-specific second loop of uL4 (yellow) remodels the tunnel cross-section, contributing to increased asphericity. Ribosomal proteins are colored: uL4 (yellow) and uL22 (blue).

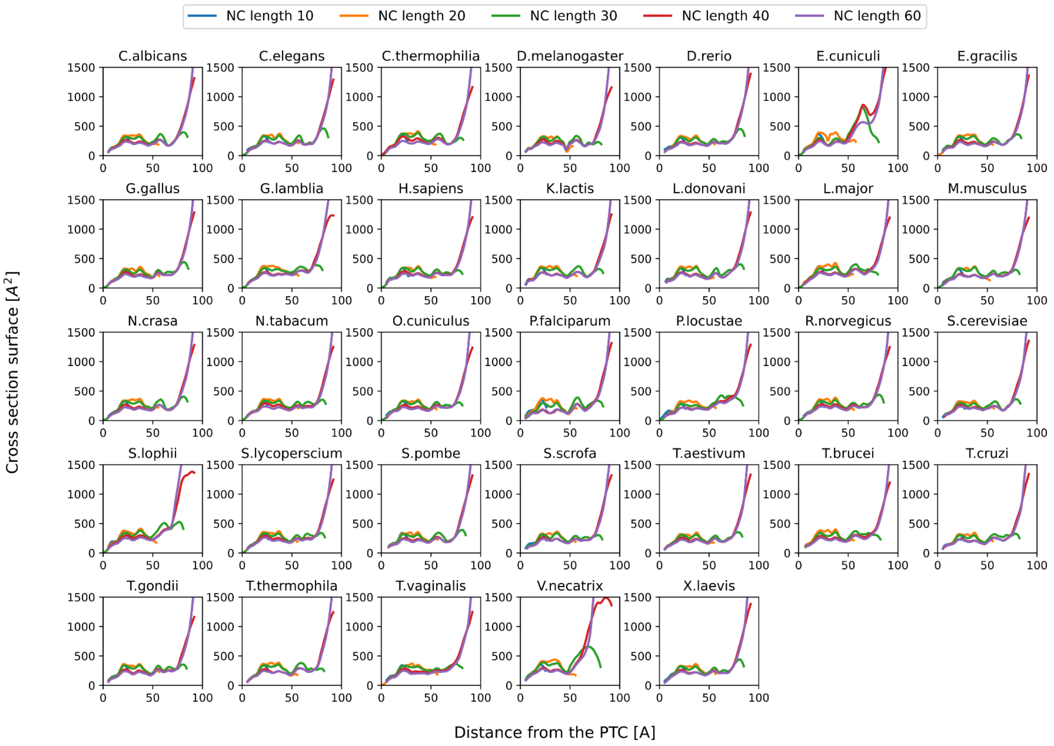


**Figure S14. Functional cross-sectional area profiles of eukaryotic ribosome exit tunnels.** Profiles were calculated for 33 eukaryotic structures using simulations with five increasing nascent chain lengths (L = 10, 20, 30, 40, and 60 residues), illustrating the length-dependent accessibility of the tunnel volume.

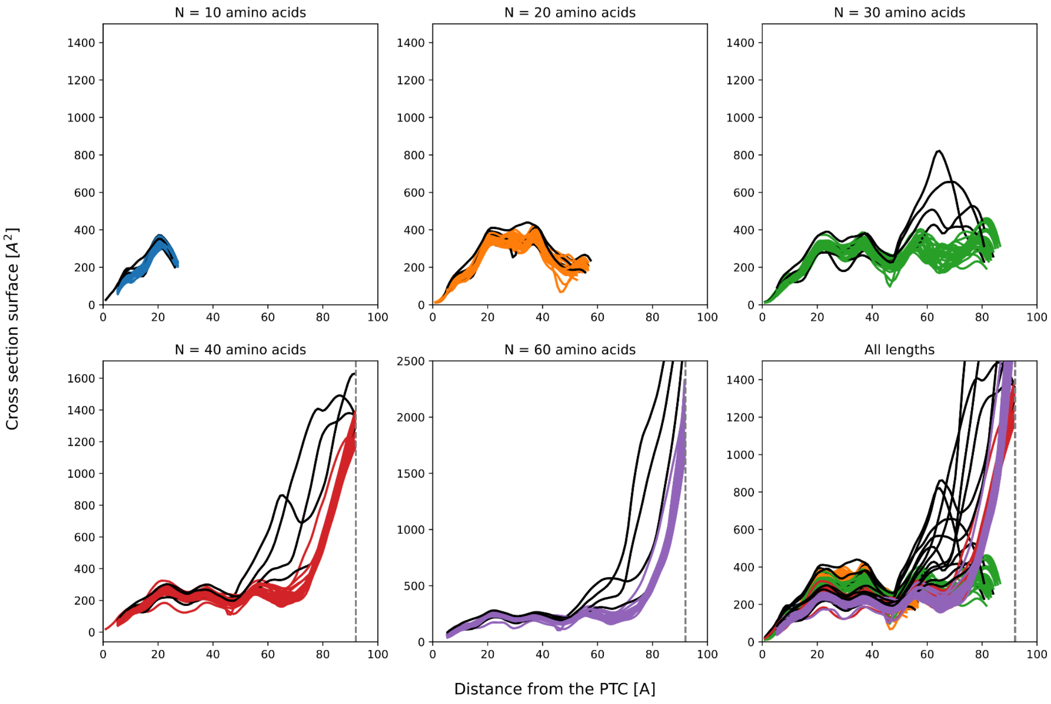

**Figure S15. Combined cross-sectional area profiles of eukaryotic ribosome tunnels.** Superimposed cross-sectional area profiles derived from all 33 eukaryotic ribosomes. Profiles corresponding to Microsporidia species are highlighted in black, illustrating their divergent tunnel architecture compared to canonical eukaryotes. The functional exit boundary is indicated by a vertical grey dotted line.

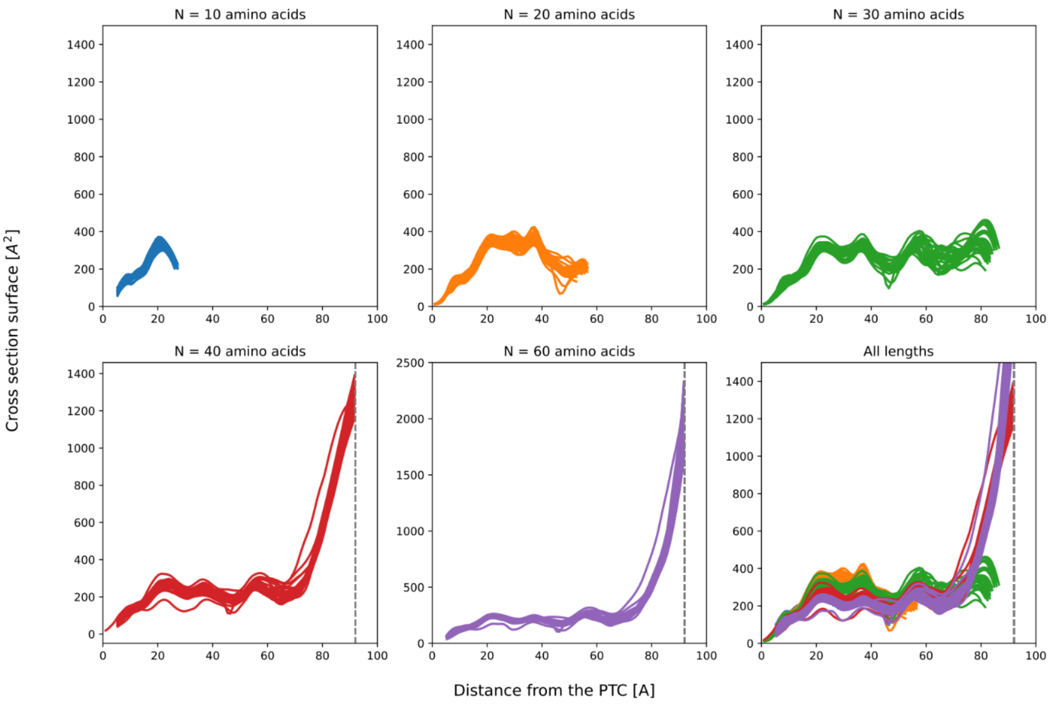


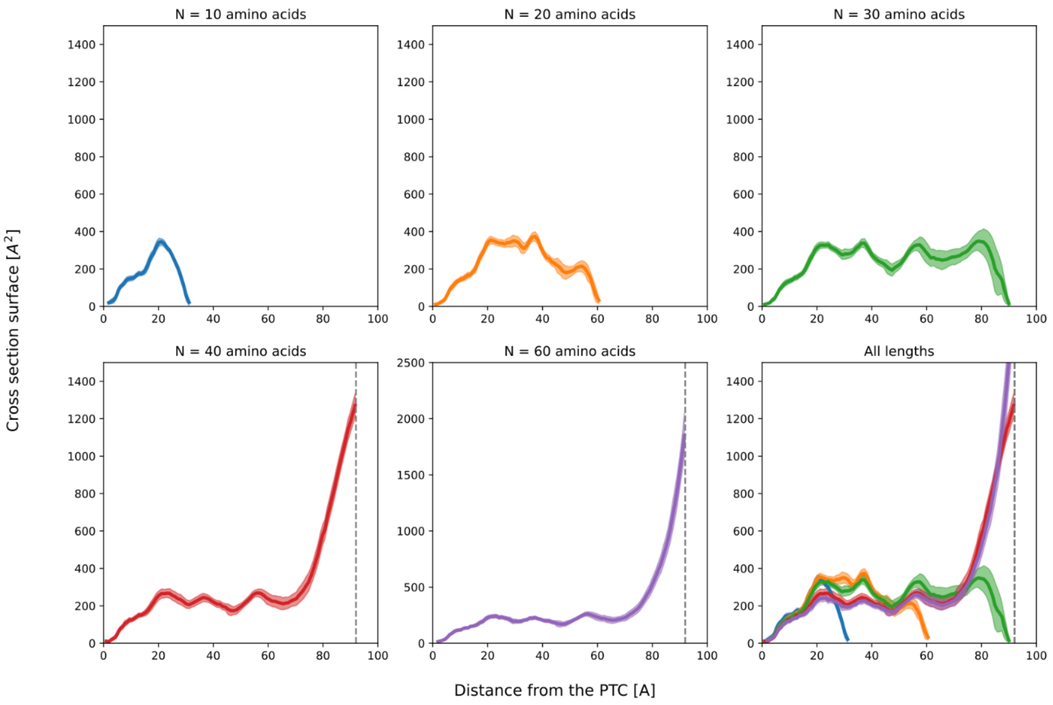

**Figure S16. Combined cross-sectional area profiles of canonical eukaryotic ribosome tunnels. (Top)** Superimposed cross-sectional area profiles derived from eukaryotic ribosomes, excluding Microsporidia structures, to visualise the structural conservation of the canonical eukaryotic tunnel. **(Bottom)** The average cross-sectional area profile for the canonical eukaryotic dataset, with the shaded region representing the standard deviation (± SD). A vertical grey dotted line indicates the functional exit boundary of the tunnel.

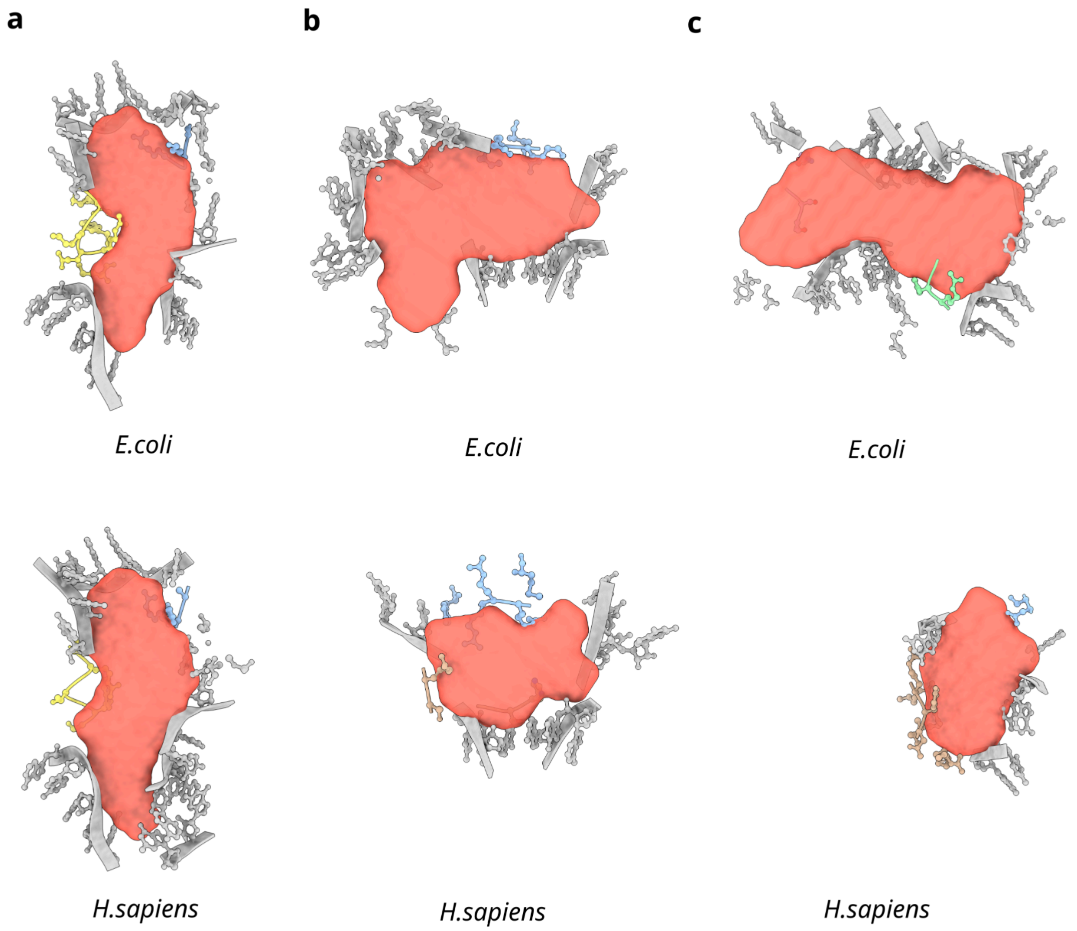


**Figure S17. Structural closure of ancestral side tunnels in the eukaryotic ribosome. (A)** Comparison of the first constriction site in *E. coli* (top) and *H. sapiens* (bottom). **(B)** The first side tunnel region: *E. coli* (top) displays an accessible lateral cavity, whereas the corresponding region in *H. sapiens* (bottom) is structurally occluded. **(C)** The second side tunnel region: The open cavity found in *E. coli* (top) is physically sealed in *H. sapiens* (bottom) by the eukaryotic-specific protein eL39 (light brown). Ribosomal proteins are colored: uL4 (yellow), uL22 (blue), uL23 (green), and eL39 (light brown).

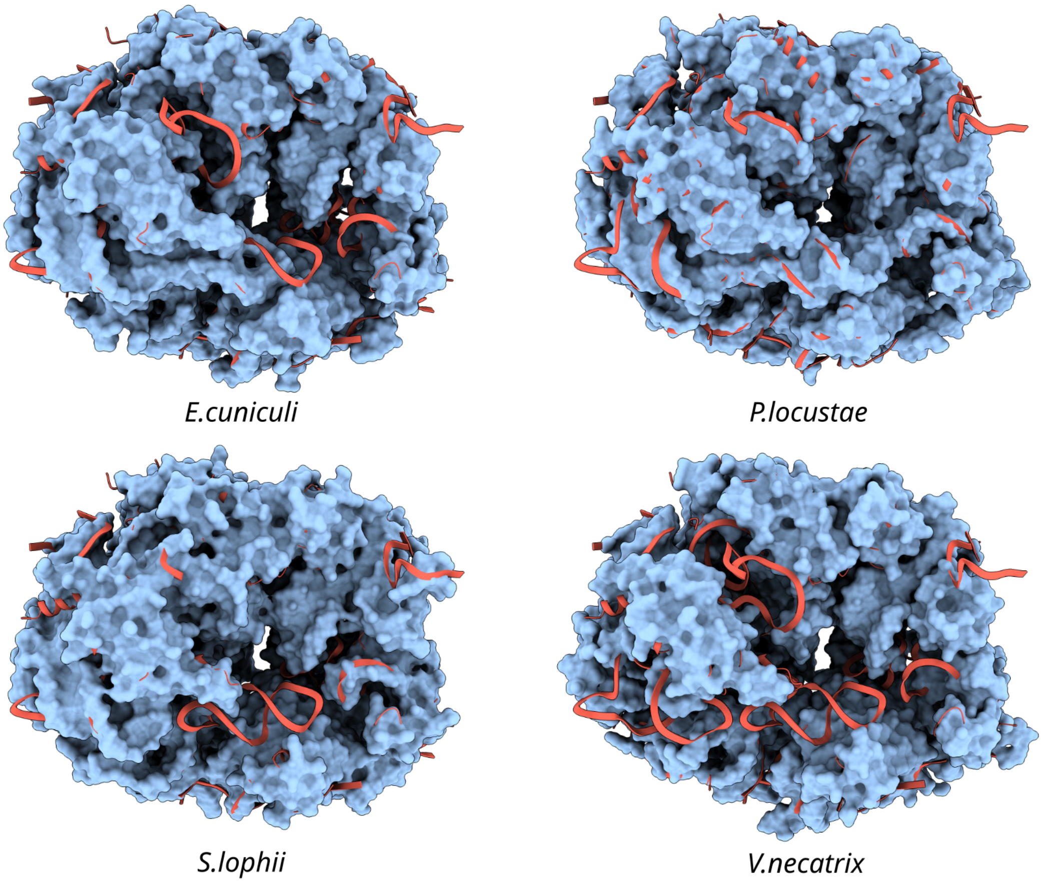


**Figure S18. Structural divergence of the Microsporidia exit tunnel.** Superposition of the Microsporidia exit tunnel volume (blue surface) onto the *H. sapiens* ribosome structure (red cartoon), viewed from the top. This comparison illustrates the distinctive reduced geometry of the parasitic tunnel relative to the canonical human architecture.

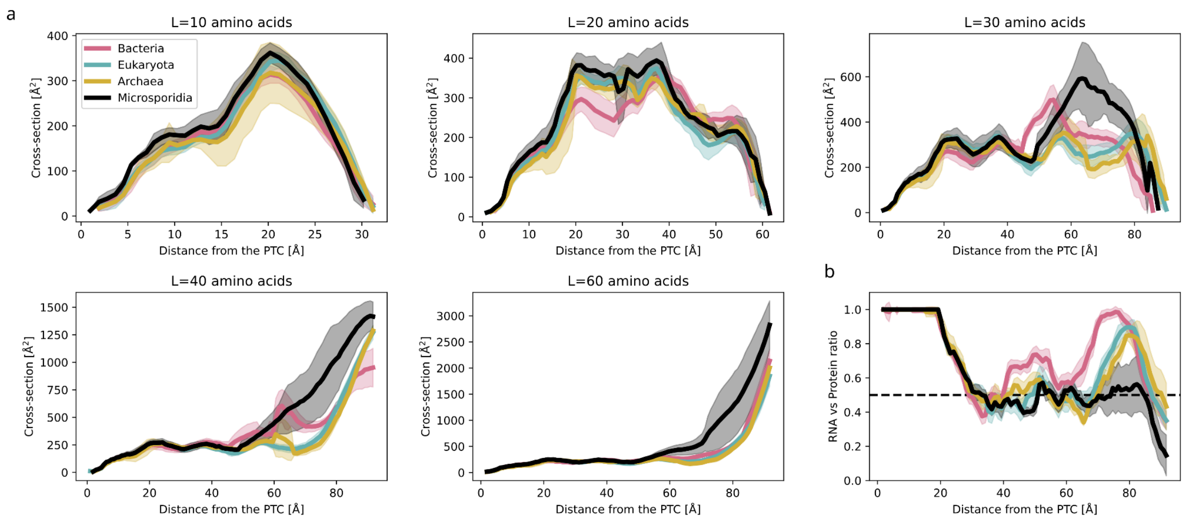


**Figure S19. Global comparison of ribosome tunnel functional profiles.** Superposition of the average cross-sectional area profiles for Bacteria (red), Archaea (orange), canonical Eukaryotes (cyan), and Microsporidia (black). The shaded regions represent the standard deviation (± SD) for each group. This comparison illustrates the distinct geometric divergence between the domains.
