## Supplementary Tables for "Evolution of the ribosomal exit tunnel through the eyes of the nascent chain"

| ID | Organism | Phylum | Type of ribosome | PDB ID | Resolution [Å] |
|---|---|---|---|---|---|
| *1* | *A.baumannii* | Pseudomonadota | P-site tRNA | 6v39 | 3.04 |
| *2* | *B.burgdorferi* | Spirochaetota | empty ribosome | 8fn2 | 3.40 |
| *3* | *B.subtilis* | Bacillota | P-site tRNA | 7as8 | 2.90 |
| *4* | *C.acnes* | Actinomycetota | P-site tRNA | 8crx | 2.78 |
| *5* | *D.radiodurans* | Deinococcota | empty ribosome | 5dm6 | 2.90 |
| *6* | *E.coli* | Pseudomonadota | P-site tRNA | 7zp8 | 2.20 |
| *7* | *E.faecalis* | Bacillota | P-site tRNA | 7p7u | 3.10 |
| *8* | *F.johnsoniae* | Bacteroidota | empty ribosome | 7jil | 2.80 |
| *9* | *L.innocua* | Bacillota | P/E-site tRNA | 8uu5 | 3.00 |
| *10* | *L.lactis* | Bacillota | empty ribosome | 5myj | 5.60 |
| *11* | *L.monocytogenes* | Bacillota | empty ribosome | 8a63 | 3.10 |
| *12* | *M.pneumoniae* | Mycoplasmatota | A- P-site tRNAs | 7pal | 4.70 |
| *13* | *M.smegmatis* | Actinomycetota | P-site tRNA | 8fr8 | 2.76 |
| *14* | *M.tuberculosis* | Actinomycetota | P- E-site tRNAs | 7mt7 | 2.71 |
| *15* | *P.aeruginosa* | Pseudomonadota | empty ribosome | 6spd | 3.28 |
| *16* | *P.urativorans* | Pseudomonadota | P-site tRNA | 8rdv | 2.60 |
| *17* | *S.aureus* | Bacillota | empty ribosome | 6hma | 2.65 |
| *18* | *T.thermophilus* | Deinococcota | A- P- E-site tRNAs | 8cvl | 2.30 |

**Table S1**. List of bacterial ribosome structures selected from the PDB search.

| ID | Organism | Phylum | Type of ribosome | PDB ID | Resolution [Å] |
|---|---|---|---|---|---|
| *1* | *C.albicans* | Ascomycota | empty ribosome | 7pzy | 2.32 |
| *2* | *C.elegans* | Nematoda | A- P-site tRNAs | 9bh5 | 2.63 |
| *3* | *C.thermophilia* | Ascomycota | P/E-site tRNA | 7old | 3.00 |
| *4* | *D.melanogaster* | Arthropoda | E-site tRNA | 4v6w | 6.00 |
| *5* | *D.rerio* | Chordata | empty ribosome | 7oyb | 2.40 |
| *6* | *E.cuniculi* | Microsporidia | empty ribosome | 7qep | 2.70 |
| *7* | *E.gracilis* | Euglenozoa | A- P- E-site tRNAs | 6zj3 | 3.15 |
| *8* | *G.gallus* | Chordata | E-site tRNA | 8q7z | 2.50 |
| *9* | *G.lamblia* | Fornicata | empty ribosome | 7pwg | 2.75 |
| *10* | *H.sapiens* | Chordata | A- P-site tRNAs | 8jdk | 2.26 |
| *11* | *K.lactis* | Ascomycota | P/E-site tRNA | 6uz7 | 3.60 |
| *12* | *L.donovani* | Euglenozoa | empty ribosome | 5t2a | 2.90 |
| *13* | *L.major* | Euglenozoa | empty ribosome | 8a3w | 2.89 |
| *14* | *M.musculus* | Chordata | E-site tRNA | 7ls2 | 3.10 |
| *15* | *N.crasa* | Ascomycota | A- P-site tRNAs | 7r81 | 2.70 |
| *16* | *N.tabacum* | Streptophyta | A/P- P/E-site tRNAs | 8b2l | 2.20 |
| *17* | *O.cuniculus* | Chordata | P-site tRNA | 6sgc | 2.80 |
| *18* | *P.falciparum* | Apicomplexa | P-site tRNA | 3jbn | 4.70 |
| *19* | *P.locustae* | Microsporidia | empty ribosome | 6zu5 | 2.90 |
| *20* | *R.norvegicus* | Chordata | A/P- P/E-site tRNAs | 7qgg | 2.86 |
| *21* | *S.cerevisiae* | Ascomycota | A/P- P/E-site tRNAs | 8ccs | 1.97 |
| *22* | *S.lophii* | Microsporidia | empty ribosome | 7qca | 2.79 |
| *23* | *S.lycoperscium* | Streptophyta | A- P-site tRNAs | 7qiz | 2.38 |
| *24* | *S.pombe* | Ascomycota | empty ribosome | 8eug | 2.80 |
| *25* | *S.scrofa* | Chordata | empty ribosome | 3j7p | 3.50 |
| *26* | *T.aestivum* | Streptophyta | P-site tRNA | 4v7e | 5.50 |
| *27* | *T.brucei* | Euglenozoa | empty ribosome | 8ova | 2.47 |
| *28* | *T.cruzi* | Euglenozoa | empty ribosome | 5t5h | 2.54 |
| *29* | *T.gondii* | Apicomplexa | empty ribosome | 5xxb | 3.17 |
| *30* | *T.thermophila* | Ciliophora | empty ribosome | 4v8p | 3.52 |
| *31* | *T.vaginalis* | Parabasalia | empty ribosome | 5xy3 | 3.20 |
| *32* | *V.necatrix* | Microsporidia | empty ribosome | 6rm3 | 3.40 |
| *33* | *X.laevis* | Chordata | empty ribosome | 7oyc | 2.40 |

**Table SI 2.** List of cytoplasmic eukaryotic ribosome structures selected from the PDB search.

| ID | Organism | Phylum | Type of ribosome | PDB ID | Resolution [Å] |
|---|---|---|---|---|---|
| *1* | *H.marismortui* | Euryarchaeota | empty ribosome | 1s72 | 2.40 |
| *2* | *P.calidifontis* | Thermoproteota | empty ribosome | 9e71 | 2.36 |
| *3* | *P.furiosus* | Euryarchaeota | P- E-site tRNAs | 4v6u | 6.60 |
| *4* | *S.acidocaldarius* | Thermoproteota | empty ribosome | 8hku | 2.72 |
| *5* | *T.kodakarensis* | Euryarchaeota | empty ribosome | 6th6 | 2.55 |

**Table SI 3.** List of archaeal ribosome structures selected from the PDB search.

| Organism | Phylum | Type of ribosome | PDB ID | Resolution [Å] |
|---|---|---|---|---|
| ***Mitochondrial*** | | | | |
| *A. thaliana / B. oleracea* | Streptophyta | empty ribosome | 6xyw | 3.86 |
| *C. reinhardtii* | Chlorophyta | empty ribosome | 7pkt | 3.00 |
| *H.sapiens* | Chordata | A- P-site tRNAs | 6zm5 | 2.89 |
| *L. tarentolae / L. major* | Euglenozoa | empty ribosome | 7aih | 3.60 |
| *N.crassa* | Ascomycota | P-site tRNA | 6ywy | 3.05 |
| *P.magna* | Chlorophyta | P-site tRNA | 8apn | 3.10 |
| *S.cerevisiae* | Ascomycota | P-site tRNA | 5mrc | 3.25 |
| *S.scrofa* | Chordata | P-site tRNA | 6ydp | 3.00 |
| *T.brucei* | Euglenozoa | empty ribosome | 6hiv | 7.80 |
| *T.gondii* | Apicomplexa | empty ribosome | 9g6k | 2.89 |
| *T. thermophila* | Ciliophora | empty ribosome | 6z1p | 3.70 |
| ***Chloroplast*** | | | | |
| *S.oleracea* | Streptophyta | empty ribosome | 5x8p | 3.40 |

**Table SI 4.** List of organellar ribosome structures identified in the PDB search.

| Organism | PDB ID | Problems with the exit tunnel region |
|---|---|---|
| *A.baumannii* | 6v39 | None |
| *B.burgdorferi* | 8fn2 | None |
| *B.subtilis* | 7as8 | None |
| *C.acnes* | 8crx | Sarecyclin bound in the exit tunnel |
| *D.radiodurans* | 5dm6 | U2585 was removed to make space for the NC |
| *E.coli* | 7zp8 | None |
| *E.faecalis* | 7p7u | None |
| *F.johnsoniae* | 7jil | uL23 and uL24 loops are missing, modelled based on the AF models; U2583 was removed to make space for NC |
| *L.innocua* | 8uu5 | None |
| *L.lactis* | 5myj | None |
| *L.monocytogenes* | 8a63 | None |
| *M.pneumoniae* | 7pal | None |
| *M.smegmatis* | 8fr8 | None |
| *M.tuberculosis* | 7mt7 | None |
| *P.aeruginosa* | 6spd | None |
| *P.urativorans* | 8rdv | None |
| *S.aureus* | 6hma | U2612 was removed to make space for the NC |
| *T.thermophilus* | 8cvl | None |

| Organism | PDB ID | Problems with the exit tunnel region |
|---|---|---|
| *C.albicans* | 7pzy | None |
| *C.elegans* | 9bh5 | None |
| *C.thermophilia* | 7old | None |
| *D.melanogaster* | 4v6w | None |
| *D.rerio* | 7oyb | None |
| *E.cuniculi* | 7qep | missing a short fragment of the rRNA helix (U1134,G1135); model of the full helix we get from AF3, and upon superposition, we model the missing two nucleotides |
| *E.gracilis* | 6zj3 | None |
| *G.gallus* | 8q7z | None |
| *G.lamblia* | 7pwg | missing a short fragment of the rRNA (U335-C337) and part of |

| | | |
|---|---|---|
| | | ribosomal protein L31e (D75-K79); modelled with AF2/3 models |
| *H.sapiens* | 8jdk | None |
| *K.lactis* | 6uz7 | None |
| *L.donovani* | 5t2a | None |
| *L.major* | 8a3w | None |
| *M.musculus* | 7ls2 | None |
| *N.crasa* | 7r81 | uL29 loop is missing (S40-L45), we modelled it using the AF2 structure |
| *N.tabacum* | 8b2l | None |
| *O.cuniculus* | 6sgc | None |
| *P.falciparum* | 3jbn | eL39 loop is missing (K30-Y36), we modelled it using the AF2 structure |
| *P.locustae* | 6zu5 | None |
| *R.norvegicus* | 7qgg | None |
| *S.cerevisiae* | 8ccs | None |
| *S.lophii* | 7qca | Modelling of the missing rRNA region (U417-A421) |
| *S.lycoperscium* | 7qiz | None |
| *S.pombe* | 8eug | None |
| *S.scrofa* | 3j7p | None |
| *T.aestivum* | 4v7e | None |
| *T.brucei* | 8ova | None |
| *T.cruzi* | 5t5h | None |
| *T.gondii* | 5xxb | uL4 loop is missing G83, we modelled it using the AF2 structure |
| *T.thermophila* | 4v8p | None |
| *T.vaginalis* | 5xy3 | None |
| *V.necatrix* | 6rm3 | Missing short fragments of rRNA: U258-U260, U1141-U1142 and A1171-U1172 were modelled with AF3. We were not able to model A65-U69, but it is further from the tunnel, so it should not affect the exit tunnel path |
| *X.laevis* | 7oyc | None |

| Organism | PDB ID | Problems with the exit tunnel region |
|---|---|---|
| *H.marismortui* | 1s72 | eL39 has missing region (E33-Q35), we modelled it using the AF2 structure |
| *P.calidifontis* | 9e71 | None |
| *P.furiosus* | 4v6u | None |
| *S.acidocaldarius* | 8hku | None |
| *T.kodakarensis* | 6th6 | None |

**Table SI 5.** List of structural modifications applied to the ribosome exit tunnel models.
